# Cell fate dynamics reconstruction identifies TPT1 and PTPRZ1 feedback loops as master regulators of differentiation in pediatric glioblastoma-immune cell networks

**DOI:** 10.1101/2023.10.03.560663

**Authors:** Abicumaran Uthamacumaran

## Abstract

Pediatric glioblastoma is a complex dynamical disease that is difficult to treat due to its multiple adaptive behaviors driven largely by phenotypic plasticity. Integrated data science and network theory pipelines offer novel approaches to study glioblastoma cell fate dynamics, particularly phenotypic transitions over time. Here we used various single-cell trajectory inference algorithms to infer signaling dynamics regulating pediatric glioblastoma-immune cell networks. We identified GATA2, PTPRZ1, TPT1, MTRNR2L1/2, OLIG1/2, SOX11, PDGFRA, EGFR, S100B, WNT, TNF*α*, and NF-kB as critical transition genes or signals regulating glioblastoma-immune network dynamics, revealing potential clinically relevant targets. Further, we reconstructed glioblastoma cell fate attractors and found complex bifurcation dynamics within glioblastoma phenotypic transitions, suggesting that a causal pattern may be driving glioblastoma evolution and cell fate decision-making. Together, our findings have implications for the development of targeted therapies against glioblastoma, and the continued integration of quantitative approaches to understand pediatric glioblastoma tumour-immune interactions.

## INTRODUCTION

The integration of single-cell multi-omics and computational algorithms has significantly improved our understanding of the complex molecular networks and processes driving cancer evolution (Guo et al., 2020; Lee et al., 2020; Hu et al., 2020; Wagner and Klein, 2020). Despite these recent advancements, glioblastoma, a deadly and aggressive brain tumor, remains an intractable problem in precision oncology with poor overall outcomes. Astrocytomas account for about 50% of primary brain tumours in children, with glioblastoma being the most lethal stage, whereas glioblastomas are the most common primary brain tumours in adults. Treatment of glioblastoma involves surgical resection of the tumour followed by combination radiotherapy, and chemotherapy with temozolomide (Louis et al., 2016; Wyss et al., 2022). The implementation of this approach as standard-of-care in glioblastoma has provided significant survival improvements however all glioblastoma patients will experience recurrences. Adaptation and resistance to treatment translates to a short median overall survival time of just 15 months (Louis et al., 2016) in adults. Recent advancements in precision oncology such as immunotherapies and multimodal targeted therapies are only showing beneficial outcomes in low-grade gliomas, and they nonetheless bring about issues related to cytotoxicity and systemic side effects leading to poor quality of life (Chatwin et al., 2021; Nguyen et al., 2022).

The poor overall survival prognosis of glioblastoma can be partially attributed to tumor cell infiltration and migration (Sanai and Berger, 2008). Glioblastoma also exhibits complex adaptive behaviors such as phenotypic plasticity, immune evasion/escape, and long-range inter-cellular communication that allow tumor ecosystems to be resilient, develop aggressivity and confer therapeutic resistance by expanding the functional and phenotypic intra-tumoral heterogeneity of cellular states, reducing the long-term efficacy of current multimodal treatments (Celiku et al., 2019; Wang et al., 2021; Uthamacumaran and Craig, 2022). These complex adaptive behaviors and cell state transitions are largely believed to be due to the dynamic interactions between the tumor microenvironment and a subset of glioblastoma cells referred to as glioblastoma-derived stem cells (GSCs) (Cheng et al., 2010; Jackson et al., 2015; Wang et al., 2021). GSCs are epigenetically plastic cells within the glioblastoma ecosystem exhibiting self-renewal and differentiation potency towards multiple molecularly and metabolically distinct phenotypes (Prager et al., 2020; Wang et al., 2021). In response to their microenvironmental interactions, GSCs control several regulator mechanisms, including tumor initiation, tumor recurrence/resilience, epigenetic heterogeneity, therapy resistance, and adaptive phenotypic transitions, within the glioblastoma ecosystem (Prager et al., 2020; Wang et al., 2021; Su et al., 2021). Given the difficulty of treating glioblastoma and its recurrence driven partially by GSC network dynamics, new approaches are needed to better understand the mechanisms of disease progression and identify new targets with clinical potential (Jackson et al., 2015; Gimple et al., 2019; Larsson et al., 2021; De Silva et al., 2023).

Complex systems are irreducible dynamical systems composed of many nonlinearly interacting parts that give rise to collective behaviours and self-organized patterns across different scales and over time (Wolfram, 1988; Bossomaier and Green, 2000; Mitchell, 2011). Such systems consist of multiscale processes and dynamic feedback loops, making traditional statistical approaches inadequate for quantifying their behavioral dynamics. Complex systems theory, or simply complexity science, advocates the pairing of mathematical modelling, machine intelligence, and algorithms with experimental data to better understand the emergent behaviors and patterns in such systems and their processes (Wolfram, 1988; Mitchell, 2011; Gros, 2011; Ladyman and Wiesner, 2020). Cancers are prime examples of these complex adaptive systems as they learn and adapt from their dynamic environments (Gros, 2011). To better understand transcriptional regulation in glioblastoma, we have previously applied complex systems theory to discriminate between cell fate decisions and transcriptional networks in adult and pediatric glioblastoma, and adult GSCs (Uthamacumaran and Craig, 2022). Previous work has also shown that the collective behaviors/dynamics of glioblastoma cell fate decisions self-organize towards causal patterns in gene expression state-space (attractors) in their developmental progression (Janson, 2012; Strogatz, 2015). Attractors correspond to causal structures characterizing the behavioral patterns of cell fate dynamics in (multiomic) gene expression state-space (Huang, 2006; Huang et al., 2009). The stability, adaptive traits, and control predictability of the cell fate dynamics are bound to these attractors, orchestrated by the complex signaling/transcriptional networks underlying glioblastoma differentiation dynamics (Uthamacumaran and Craig, 2022). Reconstructing the attractors and networks regulating glioblastoma ecosystems may help identify clinically relevant precision therapies and drug targets to control glioblastoma cellular dynamics. Further, this approach provides a robust screening method to predict the complex adaptive and causal behavioral patterns in glioblastoma systems, such as tumor progression, phenotypic plasticity, recurrence, aggressivity, and therapy resistance/response (Itik and Banks, 2010; Letellier et al., 2013; Khajanchi et al., 2018).

Our primary goal in this study was to understand how complex signaling/gene expression networks coordinate collective behaviors such as glioblastoma cell fate decision-making dynamics in pediatric glioblastoma. Given our previous results suggesting that pediatric glioblastoma is more similar to adult GSCs than adult glioblastoma (Uthamacumaran and Craig, 2022), we sought to identify transcriptional signatures controlling and regulating pediatric glioblastoma single-cell dynamics. To address this requires characterizing how the information flow across glioblastoma regulatory networks orchestrates glioblastoma cell fate behaviors. Thus, we applied a pipeline of complexity theory approaches integrated with computational physics and systems biology to single cell RNA-seq data from pediatric glioblastoma to identify key molecular regulators of the complex networks and predict their cell fate specifications/transitions. Using Waddington landscape reconstruction and network inference algorithms, we investigated glioblastoma cell fate decisions as attractor dynamics. Our analyses identified critical genes and transcription factors coordinating cell state transitions from stem-like to mature glioblastoma phenotypes as clinically targetable novel putative functional interactions orchestrating cell fate specifications in pediatric glioblastomas.

## METHODS

The general flow of our approach to reconstructing cell fate dynamics in pediatric glioblastoma is sketched out in Figure 1.

**Figure 1.**
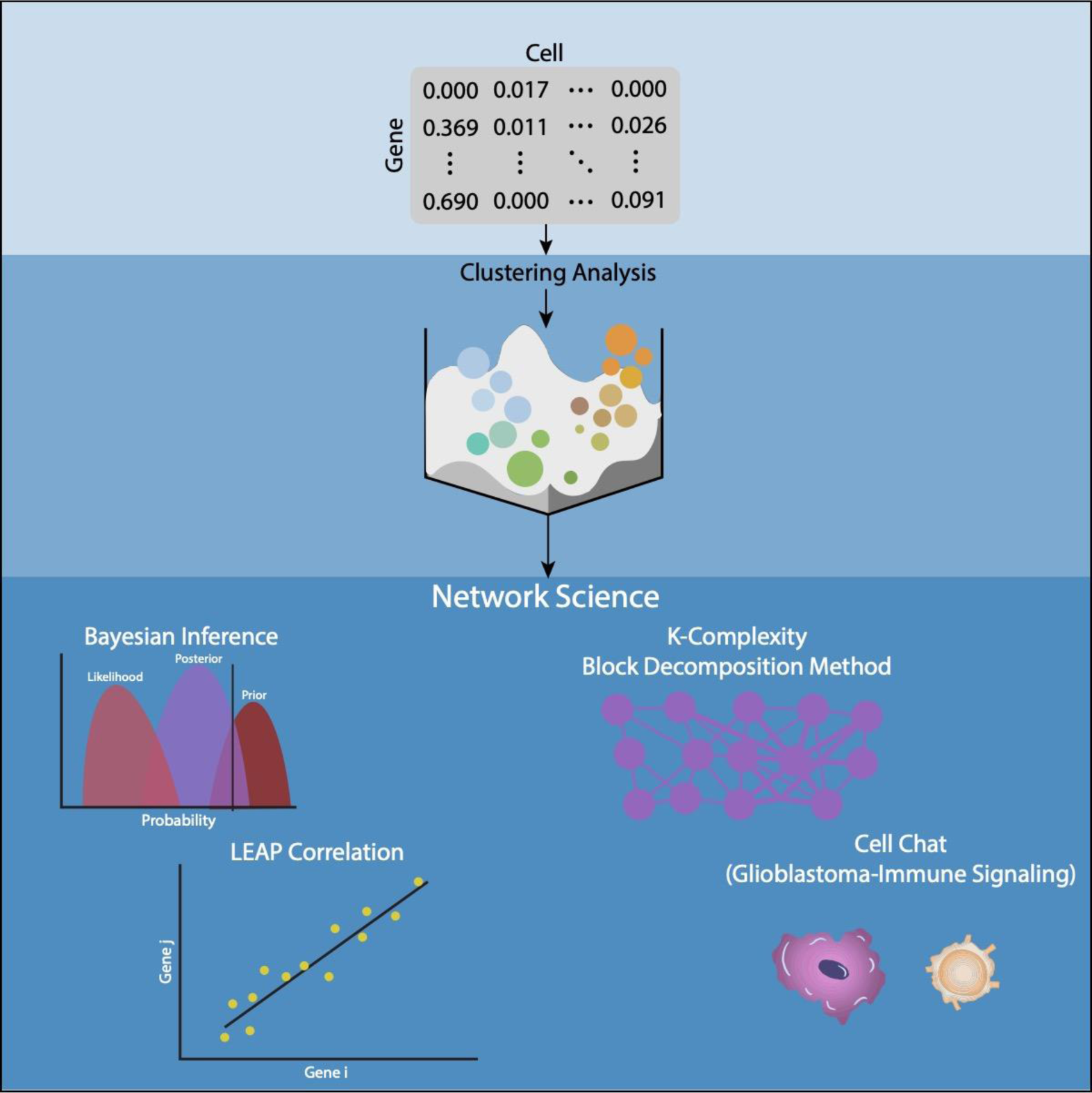
Schematic overview of pipelines/algorithms used for transcriptomic data analysis from pediatric glioblastoma single-cell RNA-Seq. Various trajectory inference/lineage tracing algorithms including clustering algorithms and Waddington attractor landscape reconstruction algorithms were employed to infer cell fate attractors, patterns in gene expression state-space to which the trajectories of cell fate differentiation dynamics are bound to. The transition genes identified by these analyses were then pooled for network science analysis. We used various statistical metrics to identify the central regulators of information flow across the differentiation networks, including Bayesian inference, maximum correlation, algorithmic complexity (Block Decomposition Method)-based network perturbation analysis, and cell-cell signaling (communication) network inference.

### Single-cell datasets

Pre-processed (i.e., filtered and log-normalized to stabilize the variance in gene expression) single-cell RNA-Seq counts of patient-derived pediatric glioblastoma cells were obtained from datasets generated by Neftel et al. (2019) from the Single Cell Portal data repository (see Data Availability and Codes). The resulting gene expression matrices consisted of the expression counts of 23,658 genes and N=1943 pediatric glioblastoma tumor cells from eight patients. These were processed as described in the protocols found in Neftel et al. (2019). Glioblastoma samples consisted of a variety of distinct cell types found in the patient-derived glioblastoma tumors, including T cells, macrophages/microglia, oligodendrocytes, and mostly malignant glioblastoma cells (Neftel et al., 2019).

### Trajectory inference

The Seurat pipeline (Stuart et al., 2018) was used as a clustering algorithm to identify differential gene expression markers in pseudotemporal space as described in Uthamacumaran and Craig (2022). Seurat was also used for clustering the network transition markers (i.e., hub genes) for the gene set enrichment (GSEA) pathway analysis (see details below).

We used various single-cell analysis trajectory inference algorithms to identify differential markers distinguishing the heterogenous phenotypic clusters within the glioblastoma population (see Supplementary Information for additional details). In particular, we applied three Waddington-landscape reconstruction algorithms to infer cell fate attractor dynamics and identify hub genes/transition markers controlling glioblastoma plasticity dynamics. Clustering and Lineage Inference in Single-Cell Transcriptional Analysis (CALISTA) was deployed to reconstruct the cell fate differentiation dynamics underlying the single-cell gene expression matrix onto a Waddington epigenetic landscape (Gao et al., 2020). Hopland was used to map successive cell fate progressions in pseudotime onto the Waddington landscape using Continuous Hopfield neural Networks (CHN), a type of recurrent artificial neural network (Guo and Zhen. 2017). Finally, MuTrans, a multiscale clustering and reduction technique for reconstructing the Waddington landscape of cell-fate dynamics (Zhou et al., 2021), was used in complement to CALISTA and Hopland. The transition genes (top ten interactions) regulating the cell state transitions towards the distinct cell fate attractors identified by these trajectory inference algorithms were extracted and pooled for network analyses. Slingshot (Street et al., 2018) and PHATE (Moon et al., 2019) algorithms were also used as validation tools for mapping cell state transitions and to further verify the robustness of the inferred attractor dynamics, or cell fate transition patterns (see results in the Supplementary Information).

Since each trajectory inference algorithm consists of a unique set of machine learning pipelines with hyperparameter dependence, the use of multiple algorithms allowed us to identify overlapping signatures that may serve as causal markers governing glioblastoma differentiation dynamics and cell state behavioral patterns. Other differentiation/transition mapping algorithms for cell fate pattern detection are presented in the Supplementary Information as validation of our findings.

### Gene regulatory network inference

The integration of different network inference methods helps to overcome the problem of causality inference in network science. Thus, we used multiple network inference algorithms to reconstruct gene regulatory networks controlling cell fate differentiation dynamics and regulating phenotypic plasticity amongst the 124 differential/transition markers identified from Seurat, Hopland, and CALISTA (Table 1).

**Table 1.** Summary of single-cell trajectory inference process. The total number of patient samples and number of single cells within each patient group remained fixed at n= 8 and N= 1943, respectively, for all clustering/trajectory inference or lineage tracing algorithms. The number of cell fate populations (phenotypic clusters) depended on the tuning of hyperparameters of the respective machine learning algorithms. A bifurcation of two differentiation pathways was observed in all cell fate trajectory inference algorithms (including those discussed in the Supplementary Information). We filtered the differential markers to top 100 most variable genes for the Seurat algorithm. The MuTrans algorithm identified more than 16 transition genes however, only markers with differential expression amidst the distinct cell identity clusters were pooled to avoid false discovery.

| Single-Cell Analysis Parameters | Seurat | CALISTA | Hopland | MuTrans |
| --- | --- | --- | --- | --- |
| Phenotypic clusters (count) | 12 | 2 | 2 | 4 |
| Cell fate trajectories (count) | 2 | 2 | 2 | 2 |
| Identified differential markers or transition genes (count) | ~100 | 14 | 10 | 16 |

We therefore combined Lag-based Expression Association for Pseudotime-series (LEAP), first-order autoregressive moving average model with a variational Bayesian Expectation-Maximization (AR1MA1-VBEM), Nonlinear Network (NLNET), and block decomposition method (BDM) as network inference metrics. LEAP is a correlation-based network reconstruction algorithm for constructing gene co-expression networks from single-cell datasets (Specht and Li, 2016). Though simple to perform, correlation analysis is not well-suited to finding causal relationships in gene regulatory networks (Specht and Li, 2016) due to the inability to identify drivers of correlated dynamics (Krieger et al., 2020). LEAP uses maximum correlation to replace the traditional Pearson’s correlation coefficient for inferring interacting gene pairs in the gene regulatory network and assessing the strength of co-expression between the pair of genes. We set the correlation metric lower cut-off threshold to be the default 0.4. The AR1MA1-VBEM framework is a Bayesian network inference algorithm for reconstructing probabilistic interaction networks (Sanchez-Castillo et al., 2018). Network performance parameters epsilon (converge criteria threshold) and delta (binary detection threshold) were tuned to 1e-10 and 0.5, respectively. NLNET is a nonlinear hierarchical clustering, and variable selection algorithm using the distance based on conditional ordered list (DCOL) metric for identifying multi-nestedness (recursiveness) and community structures (modularity) within gene expression networks (Liu et al., 2016). The random permutations parameter for inferring the gene-specific null distribution was set to n=500, and the minimum and maximum cut-off false discovery rates were set to 0.05 and 0.2, respectively. All other network inference hyperparameters/thresholds were kept to their default values.

Sixteen transition markers identified from MuTrans were analyzed independently using the partial information decomposition (PIDC) network inference algorithm described in Uthamacumaran and Craig (2022). This independent analysis was used as a blindfolded validation tool (i.e., a distinct Waddington/attractor landscape reconstruction algorithm) to determine whether matching or overlapping transition markers with our other lineage tracing algorithms will be identified by use of a complementary cell fate attractor reconstruction method. Further, the MuTrans algorithm displayed many false negatives, and hence the independent treatment ensured the robustness of our network markers as putative regulators of cell fate control and decision-making/plasticity dynamics.

Distinct network inference metrics were used to cross-validate and identify robust network signatures and patterns of gene regulation in collective cell fate dynamics/behavioral patterns. In addition, we also inferred cell-cell signaling networks using the CellChat algorithm to gain additional insights into the protein-mediated cell-cell communication pathways overlapping with the transition genes identified in the differentiation/plasticity networks. Network centrality measures were used to identify central regulators of information dynamics in the inferred glioblastoma networks.

### Network reconstruction

The topology of the gene regulatory networks was reconstructed using the Julia LightGraphs (v 1.3.) network inference package *NetworkInference.jl* (see Uthamacumaran and Craig, 2022 for further details). Network centrality measures were assessed on the LEAP and NLNET networks to study the information flow underlying the signaling dynamics using the package. Louvain and Infomap community structure detection were assessed on the inferred complex networks using the igraph R package. The centrality measures can also be computed using the network analysis features in the igraph R-package (Csardi and Nepusz, 2006).

### Inferring signaling networks

We used CellChat (Jin et al., 2021) to analyze the protein-mediated communication networks and signal transduction pathways orchestrating the glioblastoma-immune cell dynamics/interactions. CellChat is a permutation of expression-based signaling network inference tool that infers patterns of intercellular signaling pathways and communication networks from scRNA-seq data (Jin et al., 2021). The algorithm identifies cell-cell communication networks from a ligand-receptor interaction database (CellChatDB) consisting of 2021 validated putative signaling interactions using the scRNA-Seq data. Thus, it provides an indirect reconstruction tool for protein signalling networks-mediated cell-cell communication using gene expression data, whereas traditionally, proteomic datasets would be required.

### Gene set enrichment analysis (GSEA)

Two distinct gene set enrichment analysis (GSEA) algorithms were used on the transition genes/markers identified by the LEAP and Bayesian (AR1MA1-VBEM) gene regulatory networks. First, gene set analysis via the Reactome pathway database on the gene expression counts of the network markers was performed using the ReactomeGSA package (Griss, 2019). We then applied Seurat (Stuart et al., 2019) to cluster cells in the dimensionality-reduced gene-expression space, as described in Uthamacumaran and Craig (2022). Critical signaling pathways were computed by calculating the mean gene expression for every cell cluster identified by the Seurat clustering algorithm on the clustering space of the network markers. These were then analyzed for gene set variation (Uthamacumaran and Craig, 2022). Secondly, we performed a fast Wilcoxon rank sum test followed by a GSEA using the Molecular Signatures Database (MsigDB) using the presto algorithm in the R-package fgsea (Dolgalev, 2022). Two categories were selected: the oncogenic gene sets from microarray datasets of cancer gene perturbations and the hallmark gene sets to identify signaling pathways from the network markers specific to the identified cell fate clusters.

### Network perturbation analysis

We next analyzed the transcriptional networks resulting from our previous analyses to understand the patterns of network dynamics (i.e., information flow across the network) steering the glioblastoma cell fate plasticity dynamics with the goal of comparing the findings of our graph-theoretic network centrality measures to algorithmic graph complexity-based computational dynamics. This method allowed us to treat the network patterns as computational systems and study the collective cell fate decision-making dynamics as algorithmic codes (i.e., transcription programs) (Zenil et al., 2019).

In contrast to traditional statistical methods that rely heavily on linear dependencies (correlation), algorithmic complexity measures can detect causal patterns of complex dynamics and nonlinear relationships in computational system (networks) (Zenil et al., 2019). Algorithmic complexity, also known as Kolmogorov complexity or K-complexity, represents the shortest length of a computer program required to generate that system (i.e., description length), and serves as a powerful statistical metric for causal inference in network information dynamics. Thus, biologically speaking, algorithmic complexity provides a computationally robust network inference method for identifying causal relationships in gene interactions and quantify the information flow (influence) of links/nodes of the transcriptional networks in controlling/regulating cell fate behaviors.

The adjacency matrix of the Bayesian and LEAP glioblastoma networks were binarized with a cut-off threshold of 0.1 to allow computations of algorithmic complexity for the network interactions. BDM estimates of algorithmic complexity are unavailable for continuous dynamics and fuzzy systems implying that gene expression counts of the network markers need to be binarized above some threshold. The resulting unweighted Boolean graph matrix was analyzed via BDM (see above) using the online algorithmic complexity calculator’s (OACC) network perturbation analysis tool (Zenil et al., 2018). To assess which node or link had the greatest graph network complexity contribution BDM performs *in silico* shifts in the graph’s nodes and edges by deletion (knockout) (Zenil et al., 2014; Zenil et al., 2019). Virtually, the OACC allows the removal/deletion of an edge or node from the inferred graph network. The subsequent changes in the graph network complexity are provided by OACC as a table, ranking the top nodes and interactions (edges) using algorithmic complexity measures.

## RESULTS

### Waddington landscape reconstruction reveals complex attractors with a trilineage differentiation in glioblastoma cell fate transition dynamics

Cell fate decisions were inferred from the cluster plots of cell fate trajectories (i.e., differentiation pathways) using the CALISTA and Hopland trajectory inference algorithms (see Methods) to reconstruct the Waddington landscape from the single-cell transcriptomic data. We predicted a complex attractor with a single pitchfork bifurcation separating the cell lineages (Fig 2A-B) with a high degree of similarity to the patterns of cell fate clustering by inferred from transcriptional dynamics (see Figure S1 in the Supplementary information for CCAT results).

**Figure 2.**
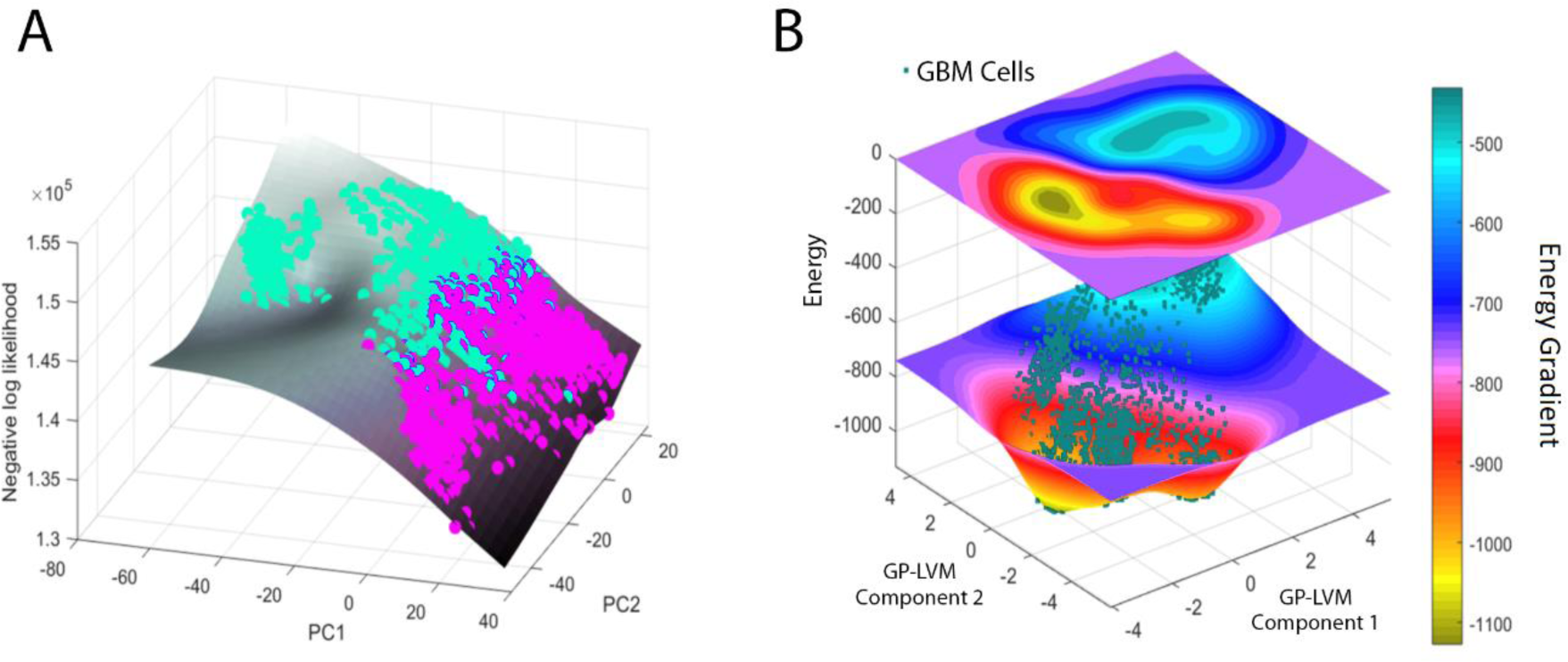
Cell fate attractor landscapes reconstruct the complex dynamics underlying glioblastoma cell fate decision-making. A) CALISTA mapped glioblastoma cell fate differentiation dynamics as a bifurcation from high energy cell states clustered higher on the attractor landscape (in turquoise) differentiating towards lower energy cell states (in pink) in principal component analysis (PCA) dimensionality reduced space. B) Glioblastoma Waddington landscape reconstructed using the Hopland algorithm. Color bar indicates the energy of the glioblastoma cells (turquoise for high energy states and yellow for low energy state attractors). The transition genes identified by the attractor reconstruction (trajectory inference) algorithms were used for network pattern analysis of distinguishing key gene markers regulating and controlling glioblastoma differentiation dynamics. A and B) Cellular phenotypes occupying higher positions on the attractor landscapes correspond to high energy cellular states with increased differentiation potency (i.e., stem cell-like). Cellular states occupying lower regions of the landscape correspond to low energy states indicating mature, well-differentiated phenotypes or transition states (intermediates) with hybrid/mixed phenotypes.

### Key transition genes and signaling pathways steering glioblastoma cell fate decisions are identified using complex systems approaches

Based on the cell fate attractor landscapes reconstructed using CALISTA and Hopland, we next identified critical transition genes regulating glioblastoma epigenetic plasticity and cell fate transitions (Fig 3A). We found that lactate dehydrogenase B (LDHB) was upregulated as glioblastoma phenotypes differentiate from higher to lower energy cellular identities. In contrast, ACTG1, NACA, TPT1, GNB2L1, and EEF1G were found to be down-regulated as glioblastoma cells transition from high (i.e., stem-cell like) to low energy cell fates (i.e., mature/differentiated phenotypes). Furthermore, we found genes involved in bioelectric fields generation and energy metabolism, such as mitochondrial NDUFA2 (a subunit of ubiquinone), LDHB (lactate dehydrogenase B), CHCHD2 (mediator of oxidative phosphorylation (OXPHOS), and ATP5A1 (subunit of mitochondrial ATP synthase) as drivers of phenotypic transitions (Fig 3A). These results suggest that bioelectric networks of the mitochondrial-electron transport chain may be key regulators of phenotypic plasticity in glioblastoma, giving rise to e.g., metabolic phenotypes (glycolytic versus OXPHOS) (Jia et al., 2018; Payne et al., 2019; Libby et al., 2020).

**Figure 3.**
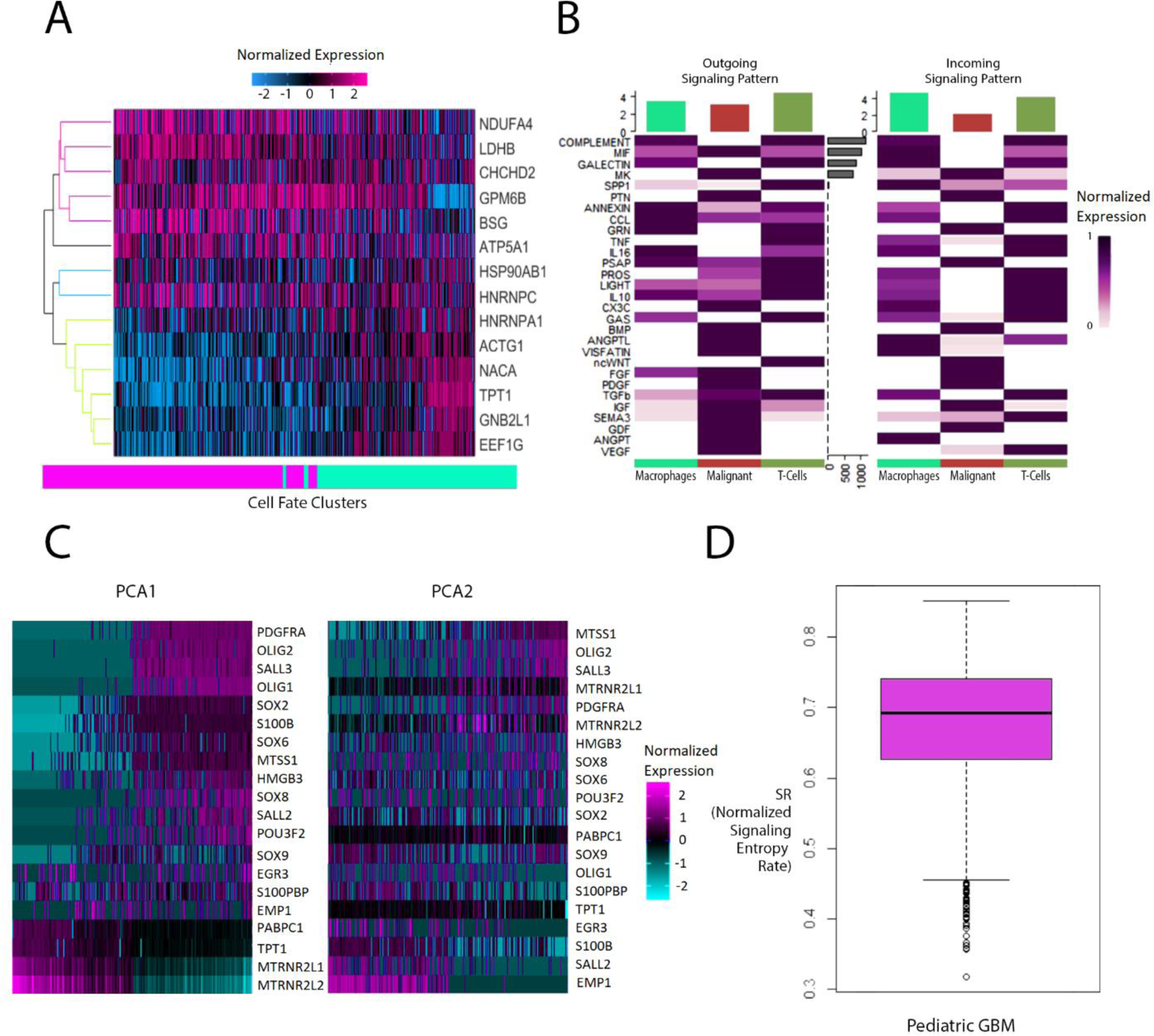
Signature expression maps of glioblastoma intracellular signaling patterns steering glioblastoma cell fate choices. A) Heatmap of transition gene signatures steering glioblastoma cell fate differentiation from low (pink clusters) energy states to high energy states (turquoise clusters) on the CALISTA attractor landscape (Figure 2A). The low and high energy cell states correspond, respectively, to stable, differentiated phenotypes and unstable, stem cell-like phenotypes. B) Signaling pathway expression patterns within the three distinct cell phenotypes inferred by the CellChat algorithm. Color bar shows a gradient of normalized expression from white (null) to dark purple (maximal). C) Heatmap signature of PCA loadings of the top differentially expressed markers identified by Seurat cell clustering and Waddington landscape algorithms. D) Differentiation potency of glioblastoma cells based on the normalized signaling entropy rate.

We also identified key cell-cell communication pathways between the three distinct phenotypes found in the glioblastoma patient tumor samples using CellChat (Fig 3B) and found that certain signaling pathways (i.e., the complement system, galectin, annexin, GRN, and PSAP) were similarly expressed in macrophages and T cells but were absent from signaling in the glioblastoma malignant cells. Growth factors such as PDGF, IGF, GDF, and PTN were strongly expressed in only the malignant tumor cells in both the incoming and outgoing signaling networks. A table of the signaling proteins with brief descriptions of their functional roles in complex cellular processes and the top PCA markers for T cells subset is provided in the Supplementary Information (Table S1 and Figure S5).

The gene expression patterns of the most differentially expressed genes pooled from the highest PCA loadings in glioblastoma phenotypic clustering were defined using Seurat and other attractor landscape algorithms (Figure 2). PDGFRA, OLIG1/2, SOX6/8, SALL2/3, POU3F2, S100B, and MTSS1 were all found to have similar expression profiles in glioblastoma cell fate transitions, whereas PABPC1, TPT1, and MTRNR2L1/2 had reverse expression patterns in the PCA loadings (Fig 3C). Although S100B is a prominent astrocyte marker, the regulatory interactions with other critical network markers are of interest to the glioblastoma’s phenotypic plasticity and cell state transitions.

We then computed the normalized signaling entropy rate (SR) for the pediatric glioblastoma cells using CCAT (Fig 3D). In cellular differentiation, SR provides a measure of the energy potential (i.e., height) of the Waddington’s epigenetic landscape and thus the phenotypic plasticity of cancer cells (Creixell et al., 2012; Teschendorff et al., 2014). Higher signaling entropy denotes stem-like cell states (i.e., higher differentiation potency), while lower entropy corresponds to mature phenotypes. We found the median SR of glioblastoma cells to be 0.69, and the mean SR to be 0.65±0.20, implying that a range of differentiation states were observed within the ensemble of glioblastoma cells.

### Identification of driver genes and signaling patterns regulating glioblastoma cell fate dynamics via network inference

The top differentially expressed markers identified in the two highest PCA loadings of the Seurat clustering and various attractor reconstruction algorithms were pooled to perform network inference analyses (Fig. 3C). In our previous study, we exploited partial information decomposition (PID), a multivariate information theoretic to quantify gene expression and transcription factor networks from glioblastoma single-cell analysis (Uthamacumaran and Craig, 2022). In this study, Pearson correlation (maximum correlation) and Bayesian inference were employed to infer higher-order relationships and multi-scaled dynamics in the gene markers, to contrast with our previously established PID glioblastoma networks. The LEAP network was used in complement to the AR1MA1-VBEM Bayesian network algorithm to infer complex networks steering cell fate decision-making and plasticity/transition dynamics within the heterogenous glioblastoma ecosystem.

Our results show that both the statistical correlation metric and Bayesian network inference tools (i.e., independent, and orthogonal network inference algorithms) reproduced the key differential biomarkers regulating glioblastoma cell fate differentiation dynamics within the clustering space that were identified in our previous study using PID networks (Uthamacumaran and Craig, 2022). Further, to understand glioblastoma-immune cell dynamics, we reconstructed signaling networks and pathways via CellChat and enrichment analysis and found that many embryonic morphogen gradients such as ncWNT, FGF, TGFβ and PDGF and neurodevelopmental signaling pathways such as BMP, CX3C and IGF are differentially expressed by the glioblastoma-immune cells. We also identified communication signals conferring immune evasion in glioblastoma systems. Other signals such as PTN and PSAP serve as neuromodulators and in neuro-glial protection. Further, PTN interaction with PTPRZ1 is critical for neural precursor cells-based signaling for glioma invasion of the subventricular zone (SVZ) (Qin et al., 2017). These SVZ neural stem cells communication pathways are critical for the origin of gliomas (Zhang et al., 2021).

From these inferred networks, we assessed key metrics to identify the nodes (genes) acting as the critical regulators of the network’s information flow and complex signalling dynamics. These included centrality measures such as betweenness (to identify the gatekeeper of information transfer in a complex network), closeness, and eigenvector centralities (Jin et al., 2021). Similarly, to infer intercellular signaling networks, the centrality metrics from graph theory used in social network analyses were used as in-built functions in the CellChat algorithm to identify the incoming and outgoing signals coordinating glioblastoma cell fate behavioral dynamics (Jin et al., 2021). Additionally, we used community structure detection algorithms such as NLNET, and Louvain community detection (see Methods) to assess the modularity and multinestedness (hubs) within the complex networks. Given that oligodendrocytes were low in cell count within the Neftel et al. (2019) gene expression matrix, we removed them from the CellChat communication network analysis.

GATA2 showed the highest betweenness (0.517), closeness (5.34), and PageRank (0.129) centralities in the Bayesian glioblastoma network (Fig 4A), whereas MTRNR2L2 had the highest eigenvector centrality (i.e., the hub score or prestige/authority score) with a score of 0.611. Both genes also had the highest degree centralities of 0.5. The highest Bayesian interaction score was found to be TPT1, which connects to both MTRNR2L2 and to GATA2 via MTRNR2L1. The interaction score of TPT1 was 0.753 for its positive feedback with MTRNR2L2 and −0.642 for its negative feedback interaction inferred with MTRNR2L1. Hence, these results suggest that TPT1 is a master regulator of the information flow across the glioblastoma network.

**Figure 4.**
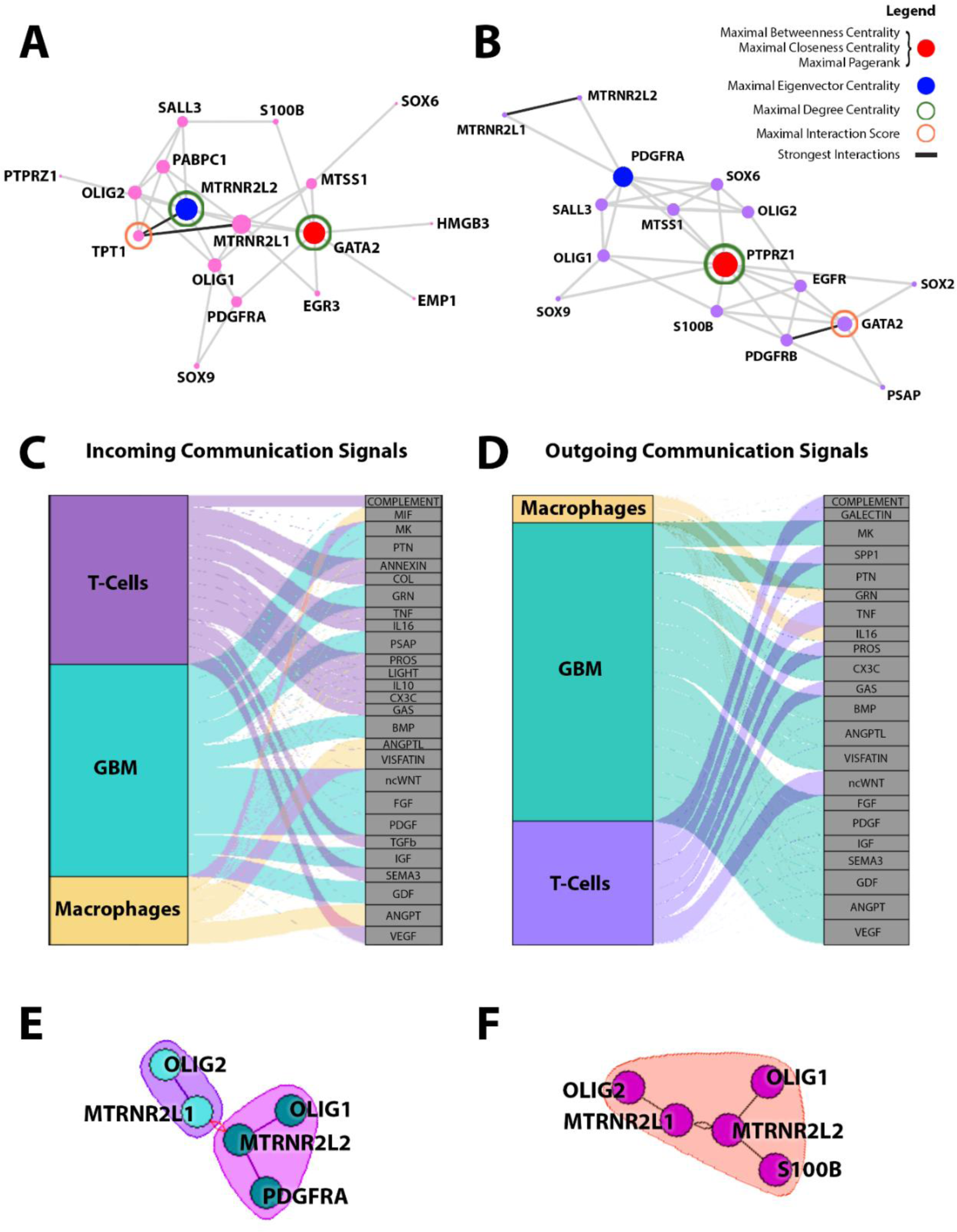
Key regulatory genes and signaling proteins driving glioblastoma cell fate decisions. A) Glioblastoma regulatory network inferred by the AR1MA1-VBEM Bayesian inference algorithm. B) Glioblastoma gene regulatory network inferred by the LEAP algorithm. A-B) Nodes represent gene markers and edges their inferred weights/scores (metrics characterizing the relationships). The Bayesian network (A) identified potential causally related probabilistic interactions between the gene markers, wherein the connectivity/topology of the network describes the information spreading. The LEAP network (B) was used as a complementary technique to infer complex networks steering cell fate decision-making within the heterogenous glioblastoma ecosystem. Node sizes of both weighted undirected networks are proportional to their degree centrality. The legend shows maximal network centrality measures used in social network analysis for identifying key genes (nodes) regulating information flow within the complex networks. Identified influential nodes corresponds to the gene marker with maximal betweenness, closeness, and PageRank centralities (red) and the gene with the maximal eigenvector centrality (blue). Green ring identifies the maximal degree centrality, and the orange the gene with the top interaction score. Darkened edges correspond to the strongest interactions. C) Incoming signaling pathways in glioblastoma cell-cell communication networks identified by CellChat algorithm. D) Outgoing signaling pathways in glioblastoma networks identified by CellChat. E) Graph Clustering label propagation community detection in glioblastoma networks using NLNET algorithm. F) Graph clustering multilevel community detection in glioblastoma network using NLNET algorithm.

In the LEAP network, PTPRZ1 was identified as the node with the highest betweenness (0.543), closeness (1.393), PageRank (0.125), and degree centralities (0.667). PDGFRA was determined to have the highest eigenvector centrality with a score of 0.446. We also identified a transcriptional feedback circuit between TPT1 and PABPC1 with an interaction score of 0.40, but it was disconnected from this graph. Similar to the Bayesian glioblastoma network analysis, GATA2 had the highest interaction score of 0.689 with PDGFRB. This suggests PTPRZ1 may be a central regulator of information flow in the complex signaling dynamics steering glioblastoma cell fate decision making.

We then applied the community detection algorithm in the NLNET network inference method and identified strong interactions between MTRNR2L1/2 and the neuronal differentiation transcription factors OLIG1/2 (Fig 4 E-F). PDGFRA and S100B were also found to be linked to these interactions, which further supports the results of the Bayesian and LEAP networks discussed above. We predicted that if certain transition genes within the differentiation networks were causally related, they would form distinct modules embedded within the network. The Louvain and Infomap community structure detection algorithms using the igraph R-package (Csardi and Nepusz, 2006) also identified three distinct regulatory network modules in both the Bayesian and LEAP networks (see Figure S4 in the Supplementary Information).

### CellChat centrality measures identify communication patterns in glioblastoma-immune cell network dynamics

To identify the signaling networks driving glioblastoma phenotypic plasticity and cell-cell crosstalk within the tumor microenvironment (TME), we used CellChat and found that immune cells in the TME secreted cytokines and chemokines mediating immune-inflammatory/wound-healing pathways, including interleukins (IL-10 and IL-16), TNF, TGF*β*, and VEGF. These signalling proteins, required to maintain/regulate the CSC niche, epithelial-mesenchymal transition (EMT) switches, and angiogenesis, were found to be communicators for the glioblastoma-associated T cells as incoming signals (Figure 4C-D). Conversely, ANGPT, VISFATIN, MIF and ANGPTL, which are associated to adipocyte metabolism, were determined to be incoming signals of glioblastoma-associated macrophages (Plaks et al., 2015; Batlle and Clevers, 2017; Su et al., 2021). Glioblastoma cells were found to receive incoming signals from all of the morphogens and growth factors required to sustain their growth processes such as GRN, PTN, MK, MIF, PSAP, BMP, non-canonical WNT, FGF, PDGF, IGF, and GDF (Fig 4 C-D). The bulk of these growth factors/morphogens and cytokines were also identified as outgoing signals from the glioblastoma cells. However, as seen in Figure 4C-D, T cells were found to be required for TNF*α* signaling in both incoming and outgoing communication networks, and as outgoing signals to the glioblastoma tumor microenvironment in the activation of the ncWNT pathway. Further, signals such as TGF*β* and WNT were determined to have a regulatory effect of critical drivers of glioblastoma stemness and plasticity dynamics (i.e., phenotypic switching). For example, tumor-associated macrophages/microglia can secrete PTN (pleiotrophin) to promote PTPRZ1 signaling in glioblastoma-stem cells to promote its adaptive properties (Shi et al., 2017). Further, high grade glioma-associated microglia were shown to be activated by TGF-b to promote the inflammasome-mediated pro-inflammatory-cytokine tumor microenvironment and thereby tumor progression (Liu et al., 2021).

### Downstream analysis identifies transition genes regulating glioblastoma cell fate differentiation and cytoskeletal rearrangement dynamics

We extended our analyses and performed cell fate landscape reconstruction using MuTrans to extract the critical transition genes predicted in the glioblastoma differentiation dynamics. The phenotypic clusters (transcriptional states) were identified in the t-SNE pattern space of the hyperparameter optimized k=4 clusters. The 2D projection of the transcriptional cell fate attractor on the dimensionality reduced gene-expression space is shown in Fig 5A, with the lineage tracing on some mean distance metric defined by the gene expression counts by the attractor landscape inference algorithm in Figure 5B. We found a distinct population of high energy cell states at the apex of the attractor landscape, corresponding to less differentiated, stem cell-like cell fates (Figure 5C). There, three differentiated phenotypes with transient cell fate flows interconnecting were determined by transcriptional similarity to have lower energy cellular states. The list of top 10 transition genes for the differentiation in between the four distinct clusters are provided in the Supplementary Information (see Figure S6).

**Figure 5.**
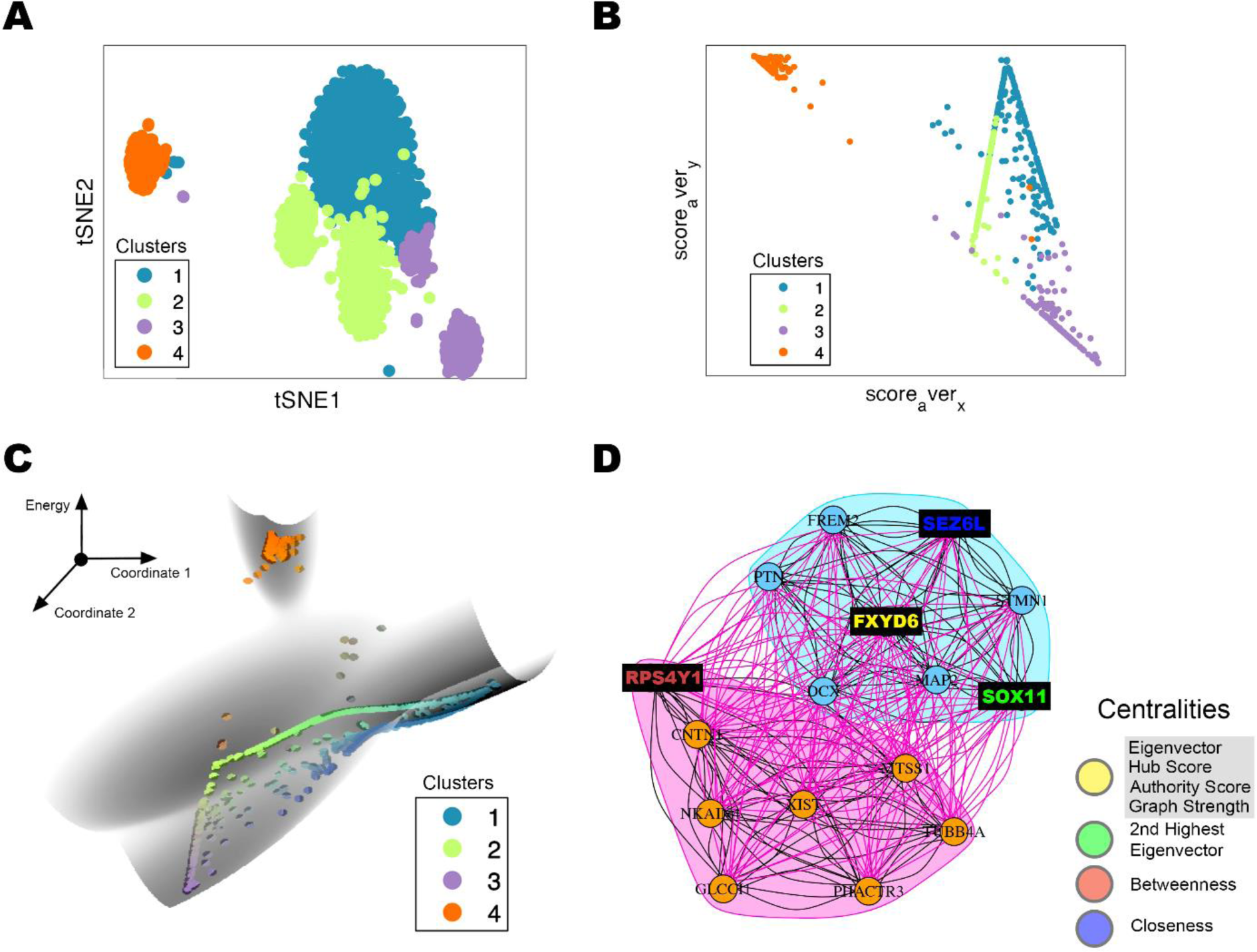
MuTrans identifies critical transition genes steering glioblastoma cell fate determinations. A) t-SNE pattern space of the assigned k=4 clusters. B) 2D-Contour of the MuTrans cell fate attractor landscape. The coordinates correspond to a two-dimensional representation the MuTrans algorithm computes based on the 2D-center position of each cell (x^2D^) with respect to center of each cell fate cluster by taking the average of x^2D^ over cells within the k-cluster membership. These average cluster scores correspond to the two coordinates. C) Two lower axes: cell fate attractor landscape obtained via a Gaussian mixture model within MuTrans with coordinates as defined in B); vertical axis: energy (φ). The energy of each cell on the landscape was determined by the negative natural logarithm of the probability distribution. As such, cell fate cluster 4 (in orange) corresponds to the highest stem-like cell fate at the top of the landscape, while the other three clusters correspond to more differentiated transcriptional states. D) Partial information decomposition network reconstructed from the top 16 transition genes identified from the landscape. The nodes corresponding to the top network centralities are highlighted by the distinct color codes.

The transition genes identified in the glioblastoma differentiation dynamics across the MuTrans cell fate landscape were analyzed using the PID algorithm, as in our previous work (Uthamacumaran and Craig, 2022). The inferred PID network was reverse-engineered using the igraph R-package and various network centrality measures were assessed using igraph network analysis functions. FXYD6 and SOX11 had the top two eigenvector centrality, hub-score, and authority-score measures. RPS4Y1, a 40S chaperone subunit, and FXYD6 had the highest betweenness scores, while SEZ6L had the highest closeness score. However, as many nodes share a similar closeness value, we determined it was not as robust of a measure as the others. gProfiler gene set enrichment analysis identified these MuTrans central network markers to be involved in neurogenesis and microtubule polymerization dynamics (see Figure S7 in Supplementary Information).

### cAMP, MYC, MTOR and NF-kB are critical signaling pathways associated with the glioblastoma driver genes

To identify the putative functional relationships between the top differentially expressed gene markers outlined in our network analyses, we next applied gene set enrichment analysis (GSEA). Cancer cell division-related pathways showed significant normalized enrichment scores, with cAMP activated signaling cascades having the greatest score followed by ESC_V6.5_UP_EARLY.V1_DN (Figure 6A). This indicates that genes that are downregulated during the early stages of differentiation of embryoid bodies from embryonic stem cells are upregulated in glioblastoma cells. MYC and MTOR were highly upregulated in the GSEA analysis, suggesting growth and cancer stemness-related pathways in glioblastoma transcriptional dynamics. Together, these findings suggest that developmental genes (and their protein products) that are epigenetically silenced in differentiated/mature cell phenotypes are reactivated in glioblastoma cells, allowing them to acquire an ESC-like stemness and phenotypic plasticity. Essential regulators of cell cycle dynamics signalling proteins such as EGFR, RAF, TBK1, MEK, P53 and cyclin D1 were also identified as significant contributors to the inferred glioblastoma network patterns/dynamics. We found 13 distinct phenotypic clusters within the seven glioblastoma patient samples (Figure 4B), indicating intratumoral heterogeneity at the level of gene expression (molecular) patterns.

**Figure 6.**
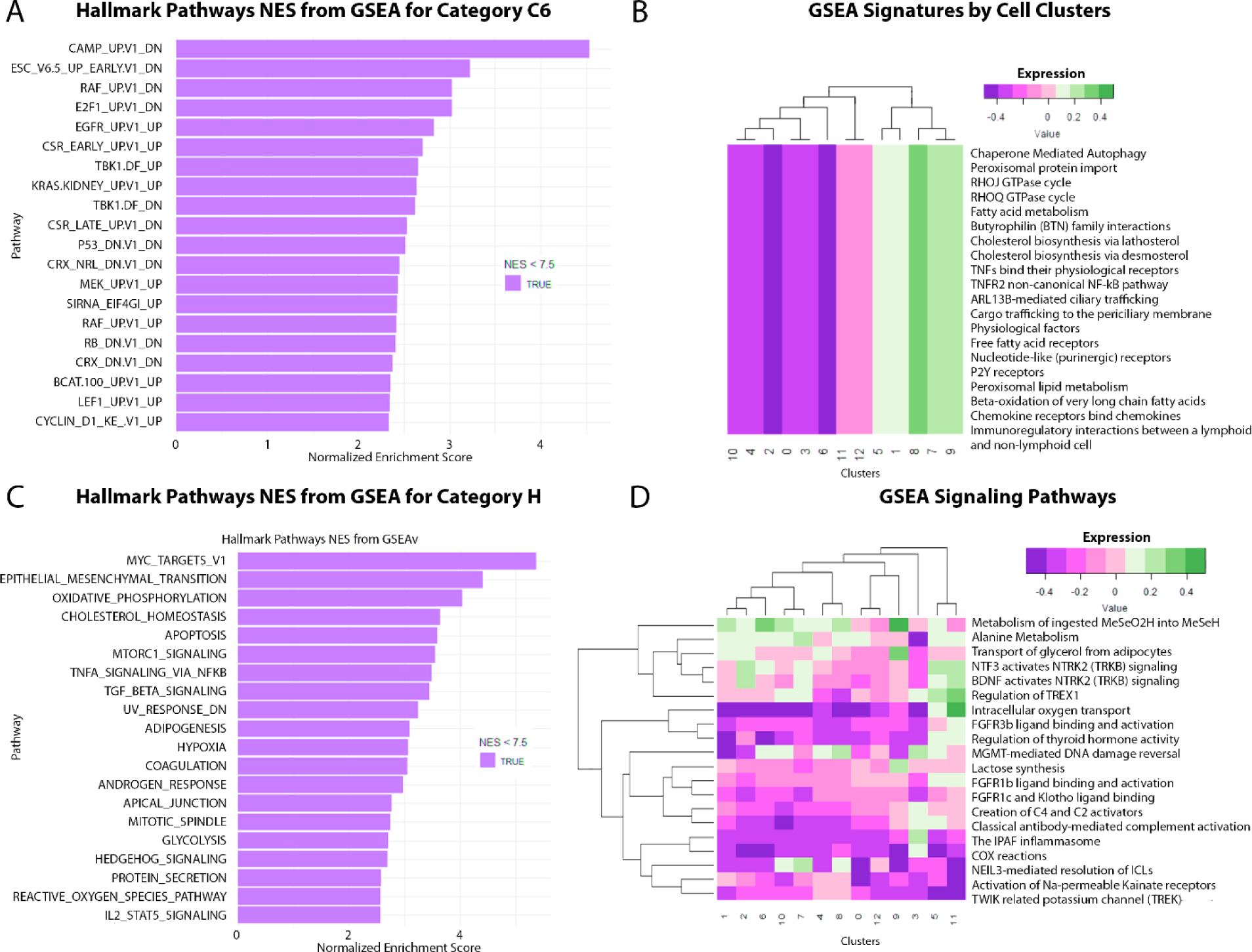
GSEA captures key signaling pathways steering glioblastoma cell fate dynamics. A) Gene enrichment signaling pathways identified for glioblastoma networks within the C6 category. B) Key signaling pathways’ expression levels classified by phenotypic cell clusters within glioblastoma. C) Key signaling pathways identified within glioblastoma networks in the H category. D) Hallmark signaling pathways associated with glioblastoma networks and their functional relationships. The color bar for figures 4B and 4D show a gradient of relative (normalized) expression levels from purple indicating down-regulation to green indicating upregulation by the indicated decimal values.

The mixed expression profiles of certain pathways were identified as metabolic pathways, which is further suggestive of the presence of the mixed, hybrid heterogeneous phenotypes within the glioblastoma population (Figure 6D). Overall, the intracellular oxygen transport had the strongest negative correlation (indicating hypoxia within the glioblastoma microenvironment), except for cluster 11 indicating the metabolic heterogeneity within these tumor ecosystems, followed by signals of the IPAF inflammasome and COX reactions. The activation of ion channels such as the TREK potassium channels and Na-permeable kainate receptors also exhibited a negative expression. These ion channels also suggest bioelectric networks regulating membrane electrophysiology may be involved in the phenotypic plasticity dynamics and morphogenesis/cellular patterning of the tumor systems. Thus, our findings propose that a targeted therapy or cocktail of these ion channels, namely kainate receptors and calcium-activated potassium channels, may induce changes in the membrane potential V_mem_ of malignant phenotypes to promote phenotypic reprogramming towards stable cell states. The strongest positive expression was observed in the MeSO_2_H metabolism and TRKB signaling amidst the distinct phenotypic clusters.

### Algorithmic complexity identifies key driver genes steering glioblastoma networks via *in silico* perturbation analyses

In our previous study, we used the block decomposition method (BDM), a measure of algorithmic complexity, as a discriminant of glioblastoma and glial stem cell (GSC) states (Uthamacumaran and Craig, 2022). Here, we used algorithmic network perturbation analysis (OACC algorithm) on the inferred glioblastoma networks to measure the shifts in BDM (and hence network complexity) as nodes and links (interactions) are knocked out (deleted) *in silico* (Figure 7). The OACC algorithm computes the algorithmic complexity of each edge and node on the graph network, by virtually deleting each of them one by one, to predict the central regulators of the network’s information dynamics. As shown in Tables 1-4, the information dynamics across these Boolean BDM networks robustly identified biomarker signatures critical for the information transfer across the regulatory networks, consistent with those identified by the network centrality measures. MTRNR2L1 and OLIG 1 were found to be the most influential nodes in maintaining the graph complexity of the Bayesian network (Table 2), as indicated by their highest perturbation rank (largest shift in BDM by node deletion), whereas the interactions between SALL3 and OLIG2 followed by those between GATA2 and S100B were found to be the most influential interactions (link perturbations with greatest BDM shift) (Table 3). In the LEAP network, PDGFRA and GATA2 had the highest node perturbation BDM scores (Table 4), whereas the associations between S100B and EGFR and OLIG1 and S100B (Table 5) were found to be the most critical interactions in establishing glioblastoma graph network complexity. A higher graph complexity implies higher network robustness/resilience and emergence of complex adaptive features. Thus, the identified nodes and links represent potential functional relationships characterizing glioblastoma transcriptional dynamics and cell fate decisions with clinical relevance (i.e., therapeutic targets).

**Figure 7.**
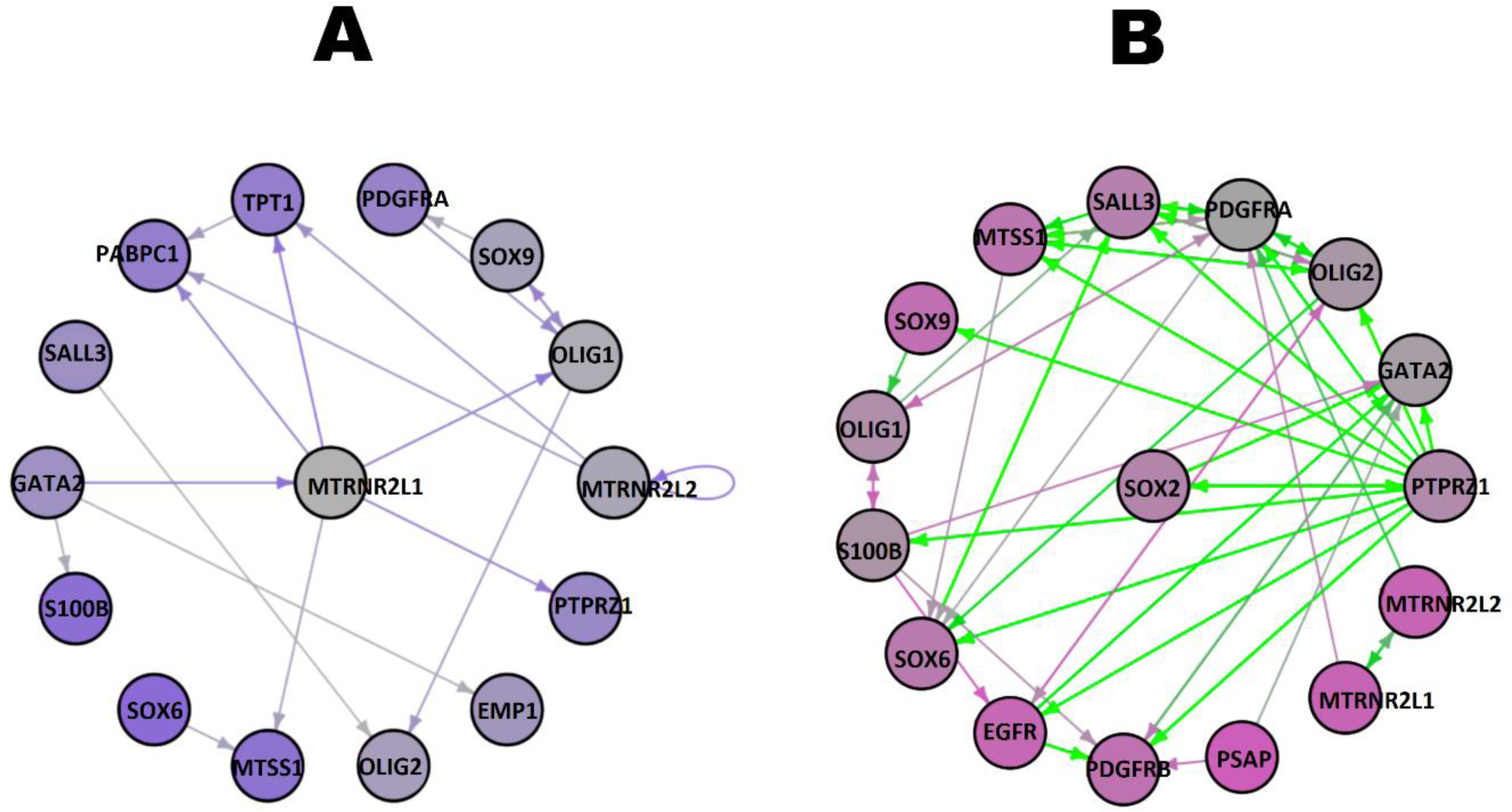
Algorithmic perturbation analysis identifies critical driver genes in glioblastoma networks using graph complexity. The algorithmic complexity measurement of unweighted, Boolean networks inferred by the binary adjacency matrix of A) Bayesian network, and B) LEAP network, as visualized using the R-implementation of online algorithmic complexity calculator (OACC). The graph network complexity was measured by the Block Decomposition Method (BDM), a robust estimate of the network’s Kolmogorov complexity, to evaluate how the graph complexity changes by node perturbations and link perturbations (see Table 2).

**Table 2.** Node perturbation analysis on graph complexity of Bayesian network. The OACC network perturbation was performed on the adjacency matrix of the Bayesian network (i.e., binarized above a threshold of 0.1) as described in the Methods. The node perturbation analysis identified the markers on the Bayesian graph network with the highest BDM shift (graph K-complexity difference when the node is deleted). The perturbation rank denotes the nodes with the highest BDM difference, which have the greatest influence on the algorithmic information dynamics of the network. As such, the node perturbations with the highest rank (closer to 1) would have the greatest causal influence on the glioblastoma network regulation (i.e., cell fate decision-making/transcriptional dynamics). Link perturbation analysis on graph complexity of Bayesian network (top 3 perturbations). Below the node perturbations, we provided the link/edge perturbation analysis of the OACC Network perturbation feature identified causally influential edges (links) of the Bayesian network by algorithmic information dynamics.

| Node | Graph BDM after Node Knockout (bits) | BDM Difference (bits) | Perturbation Rank |  |
| --- | --- | --- | --- | --- |
| MTRNR2L1 | 1104.153 | 352.554 | 1 |  |
| OLIG1 | 1167.853 | 288.854 | 2 |  |
| MTRNR2L2 | 1191.197 | 265.51 | 3 |  |
| SOX9 | 1225.578 | 231.13 | 4 |  |
| OLIG2 | 1251.67 | 205.037 | 5 |  |
| EMP1 | 1253.604 | 203.103 | 6 |  |
| GATA2 | 1268.667 | 188.041 | 7 |  |
| PTPRZ1 | 1322.649 | 133.858 | 9 |  |
| SALL3 | 1269.214 | 187.493 | 8 |  |
| PDGFRA | 1332.142 | 124.565 | 10 |  |
| TPT1 | 1347.068 | 109.64 | 11 |  |
| PABPC1 | 1347.068 | 109.64 | 11 |  |
| MTSS1 | 1349.44 | 107.268 | 13 |  |
| S100B | 1351.578 | 105.129 | 14 |  |
| SOX6 | 1402.887 | 53.82 | 15 |  |
| Source Node | Target Node | BDM after Link Knockout (bits) | BDM Difference (bits) | Perturbation Rank |
| SALL3 | OLIG2 | 1272.436 | 184.271 | 1 |
| GATA2 | S100B | 1349.577 | 146.673 | 2 |
| GATA2 | EMP1 | 1310.034 | 107.13 | 3 |

**Table 3.** Node perturbation analysis on graph complexity of LEAP Network. The node perturbation analysis from Table 2 was repeated on the LEAP network markers to identify the nodes with the highest BDM shifts (i.e., greatest causal influence on network dynamics). **Link perturbation analysis on graph complexity of LEAP network. Below the node perturbation analysis,** the link perturbation analysis performed on table 2 was repeated for the LEAP network markers to identify the relationships (links) with the highest causal influence on the network dynamics, and hence the cell fate decision-making patterns in terms of graph-network algorithmic complexity.

| Node | Graph BDM after Node Knockout (bits) | BDM Difference (bits) | Perturbation Rank |
| --- | --- | --- | --- |
| PDGFRA | 3185.863 | 876.129 | 1 |
| GATA2 | 3279.386 | 782.606 | 2 |
| OLIG2 | 3321.191 | 740.802 | 3 |
| S100B | 3329.451 | 732.542 | 4 |
| OLIG1 | 3345.36 | 716.633 | 5 |
| PTPRZ1 | 3353.902 | 708.09 | 6 |
| SOX2 | 3367.801 | 694.192 | 7 |
| SALL3 | 3493.988 | 568.004 | 8 |
| SOX6 | 3508.291 | 553.702 | 9 |
| MTSS1 | 3531.447 | 530.545 | 10 |
| PDGFRB | 3546.87 | 515.123 | 11 |
| SOX9 | 3609.438 | 454.555 | 12 |
| EGFR | 3632.472 | 429.521 | 13 |
| MTRNR2L1 | 3704.018 | 357.974 | 14 |
| MTRNR2L2 | 3704.018 | 357.974 | 14 |
| PSAP | 3719.056 | 342.937 | 16 |

**Table 3. Node perturbation analysis on graph complexity of LEAP Network.**
| Source Node | Target Node | BDM after Link Knockout (bits) | BDM Difference (bits) | Perturbation Rank |
| --- | --- | --- | --- | --- |
| S100B | EGFR | 3917.155 | 144.837 | 1 |
| OLIG1 | S100B | 3927.77 | 134.223 | 2 |
| EGFR | OLIG2 | 3934.434 | 127.559 | 3 |

## DISCUSSION

Single-cell transcriptomic analysis of the complex molecular circuitry of glioblastoma ecosystems is critical to dissect glioblastoma-immune cell networks. These methods allow us to unveil the statistical patterns and potential causal relationships within glioblastoma communication networks that may help to identify clinically relevant therapeutic targets. Thus, such analyses are a key component of the precision medicine toolbox. Here, we used complex systems approaches such as network medicine, cell fate attractor landscape reconstruction, and algorithmic perturbation analysis, to study cancer cell fate decision-making and cellular differentiation/transition processes as attractor dynamics. The complex dynamics of these processes are orchestrated by the regulatory feedback loops of cancer signaling networks. Waddington epigenetic landscape reconstruction and network medicine offer novel data-theoretic tools from systems science for causal pattern (attractors) discovery and inference of key signaling networks regulating glioblastoma cell fate decisions (Barabasi et al., 2011). In contrast to traditional bioinformatics pipelines, our systems medicine toolbox comprises of causal inference methods which detect nonlinear relationships in the network markers driving phenotypic transition dynamics.

The presence of a global pitchfork bifurcation in cell fate dynamics together with the inferred flow patterns of transient cell states indicates there may be a complex attractor steering glioblastoma cell fate choices (Fig 2A-B). Further, this bifurcation indicates a critical transition point at which cell fate determinations can change stability and cell lineage commitments. We posit that this bifurcation corresponds to the trilineage hierarchy seen in glioblastoma/pediatric high-grade glioma transcriptional dynamics as reported by Jessa et al. (2019) and Couturier et al. (2020). Generally speaking, cell fate attractors control and regulate collective cell fate behaviors/dynamics, including adaptive processes such as cell fate decision-making and phenotypic transitions, implying that attractor patterns emerge in gene expression state-space from the network dynamics of inferred transcriptional networks (Strogatz, 2015). Complex attractors with bifurcations may denote unstable cellular states or more aggressive phenotypes with difficult to treat/control adaptive behaviors including phenotypic switching/plasticity, multiscale heterogeneity, metastatic invasion, and therapy resistance (Uthamacumaran and Craig, 2022). Thus, the control of cell fate attractor dynamics by perturbation of their underlying driver networks may allow for the reprogramming of the phenotypes identified here.

The presence of a pitchfork bifurcation on the reconstructed cell fate attractor (Figure 2) indicates the presence of a causal pattern characterizing glioblastoma cell fate dynamics and the self-organization of the tumor ecosystem. In a previous study, Couturier et al. (2020) identified a trilineage hierarchy in the transcriptomic analysis of 16 adult IDH-WT glioblastoma samples. In their work, diffusion map embedding of scRNA-Seq data revealed a pitchfork-like bifurcation consisting of neuronal, astrocytic-mesenchymal, and oligodendrocytic lineages, wherein the GSC cells (enriched with specific cell surface markers) were shown to drive the phenotypic plasticity of the glioblastoma mass towards trilineage cell fates. Previously established neuronal-glioma networks driving the stemness and phenotypic plasticity in between these cell fates, such as PDGFRA, EGFR, and OLIG2, correspond with our results (Neftel et al., 2019; Wang et al., 2021; De Silva et al., 2023). Further, our findings confirm the trilineage structure is also conserved in pediatric glioblastoma. Similar conclusions were previously established in pediatric high-grade gliomas by Jessa et al. (2019), who suggested that dysregulated differentiation dynamics of hierarchical transcriptional programs resemble normal fetal neurodevelopment. Whether the glioblastoma lineage bifurcation exhibits period-doubling bifurcations, a signature of chaotic dynamics, and corresponds to a strange attractor can only be verified with time-series analysis. Importantly, the efficacy of anti-glioblastoma therapeutic interventions may be assessed by screening the robustness and stability of the inferred cell fate attractor with perturbation analysis.

Our network analyses revealed several key regulators steering glioblastoma cell fate decision-making. Most of these hub genes/driver transition markers were identified in our previous study (Uthamacumaran and Craig (2022)), and here we provide strong evidence that these are robust glioblastoma signatures controlling plasticity/differentiation dynamics given their validation through independent computational algorithms. Previous studies have shown that an increased PTPRZ1 expression is strongly related to glioblastoma stemness (Patel et al., 2014). Among the many functional targets identified by Suva et al. (2014), PTPRZ1 was one of the strongest in directly binding to SOX2, OLIG2, and POU3F2, three of the four essential transcription factors for glioblastoma stemness. PTPRZ1 has previously been reported to be highly expressed in both transcript and protein concentration within glioblastoma cells (Lu et al., 2005; Bourgonje et al., 2016). Along with GATA2 and MTRNR2L2, we found that PTPRZ1 had the highest centrality measures in our network analyses, and it was thus predicted to be a strong mediator of glioblastoma stemness. Indeed, our network analyses suggest that the feedback circuit between TPT1 and PTPRZ1 may be a master regulator of mature glioblastoma cells transitioning between CSC states and mature glioblastoma states. PTPRZ1 receptor and pleiotrophin (PTN) growth factor interaction promotes glioblastoma migration and invasion and promote neural stem cells (NSCs) signalling for glioma invasion to the subventricular zone (Lu et al., 2005; Qin et al., 2017; Zhang et al., 2021). Meanwhile, TPT1 has been shown to interact with macrophage migratory inhibitory factor (MIF) to promote the proliferation and maintenance of these NSCs, and their migration to the ventricular zone (Morimoto et al., 2022). PTN and MIF were identified as key signaling factors from the CellChat analysis, and PTN was also identified as a transition gene by MuTrans. However, unlike TPT1, the expression of PTPRZ1 was not observed in all patient cells, as seen in the PHATE glioblastoma transition maps (Figure S2). GATA2, a transcription factor critical in the differentiation of embryonic tissues into hematopoietic and central nervous systems (Lentjes et al., 2016), has been shown to promote glioblastoma progression through the EGFR/ERK pathway (Wang et al., 2013) and is closely linked to hematopoietic malignancies in children such as pediatric acute myeloid leukemia (AML) (Shiba et al., 2012).

The role of MTRNRL1/2 in glioma remain unelucidated. We identified MTRNR2L2 as a signal of PTPRZ1, suggesting its interaction in regulatory feedback loops with signals critical for glioblastoma stemness. MTRNR2L2 has previously been proposed to play a role as a neuroprotective and antiapoptotic factor (Bodzioch et al., 2009) Meanwhile, TPT1, a critical regulator of cancer stemness/plasticity is a direct target and critical regulator of the tumor suppressor gene TP53 (Chen et al., 2013). Evidence also indicates that TPT1 is a key regulator of phenotypic reprogramming of cancer cells towards cancer stem cell (CSC) fates (i.e., tumor reversion) (Amson et al., 2013). TPT1 encodes a highly conserved multifunctional protein which regulates complex cellular processes, including cell growth via the MTORC1 signaling pathway, proliferation, mitotic microtubule dynamics/stabilization, and metabolism (Bae et al., 2017; Bommer, 2017). We found that GATA2 also interacts with MTRNR2L1, while OLIG1/2 and PDGFRA, essential transcription factors mediating glioblastoma stemness and phenotypic switching, interact with MTRNR2L1/2 (Neftel et al., 2019; Wang et al., 2021). Hence, we predict that MTRNR2L1/2 genes may act as central hubs directing information flow between the master genes coordinating glioblastoma cell fate dynamics and stemness/phenotypic transitions, possibly due to its neuroprotective and anti-apoptotic role (Bodzioch et al., 2009). These findings are supported by the minimum spanning tree constructed using the Kruskal algorithm (see Figure S3 in the Supplementary Information). Many of these transition genes are essential transcription factors coordinating glioblastoma stemness (Suva et al., 2014), confirming our previously reported findings (Uthamacumaran and Craig, 2022). Furthermore, PDGFRA and EGFR, two of our central network regulators we identified here in pediatric glioblastoma, are also drivers of phenotypic plasticity in IDH-WT glioblastoma towards proneural or oligodendrocyte precursor cells (OPC) like cells, and the classical subtype or astrocytes-like cells, respectively (Verhaak et al., 2010; Neftel et al., 2019; Wang et al., 2021; De Silva et al., 2023). Meanwhile, OLIG2 is associated with OPC progenitor states and mesenchymal (MES)-like glioblastoma cell states (De Silva et al., 2023).

Amongst the high centrality nodes identified in the PID network with the MuTrans landscape transition genes, SOX11 was found to be the marker with the second highest eigenvector centrality, hub-score, and authority score, suggesting its functionally relevant importance. SOX11 is a neuronal differentiation marker and transcription factor, and hence provides a chromatin-modifying target for reprogramming glioblastoma cell fates (Su et al., 2014). Su et al. (2014) demonstrated that the synergetic lentiviral activation of SOX11 and NGN2 reprogrammed glioma cell fates towards terminal differentiation of non-proliferative neuronal phenotypes, showing that unstable glioma cell lines can be controlled by forcing them to commit towards more stable lineages. Similar approaches have been investigated in acute myeloid leukemia to reduce leukemic stem cell fractions (Laverdière et al., 2018; Da Costa et al., 2019). If verified in pediatric gliomas, such reprogramming may be clinically relevant, especially since oncohistone variants are known to be stalled in their differentiation landscape (Deshmukh et al., 2021). Nonetheless, cell fate reprogramming/forced lineage commitment experiments should be performed simultaneously on healthy controls, along with the dissection of their transcriptomic and epigenomic signatures, to help identify robust patterns which may be specific to reprogramming only the glioma cell fates. Healthy control cells should remain unaffected by such translational methods to be clinically applicable. The threshold-dependence of the various hyperparameters used in the reconstruction of the MuTrans cell fate attractor landscape must be emphasized, since increasing the value of k-clustering or the log-fold change threshold for the transition genes could change our results (here k was set to 4). Although higher number of cell fate attractors were observed with such parametric variations, we determined that the transition markers identified in our analysis (Fig 5D) were insensitive to such changes. Further, although FXYD6 had the highest eigenvector centrality/prestige score and betweenness score (along with RPS4Y1), it is a membrane ion transporter with Na+/K+-ATPase activity. It may be a coordinator of ion channel networks and hence, the bioelectric networks underlying morphological plasticity and patterning. That said, given that its inhibition could be detrimental to normal physiology, it is of little relevance to clinical drug targets. It should be noted that the transition markers from cluster 1 to cluster 3, and cluster 4 to cluster 1 from the MuTrans attractor landscape were omitted in the network analysis shown in Figure 5D since their transition strengths were found to be weak (as indicated by the color bar gradient of normalized gene expression in Figure S7). Hence, we conclude that these markers (identified through faint transitions on the heatmap) resulted from false discovery or some algorithmic noise due to their repetition in other phenotypic transitions. Other markers with weak/faint transition gradients across the pseudotemporally ordered cells, namely NCAM1, TUBB2B, LAPTM4B, and SLC35F1 (Figure S6), were also omitted from our network analyses. However, we recognize the significant role they have in cytoskeletal rearrangements such as microtubule polymerization dynamics (Figure S7).

As seen in the GSEA, the transition genes identified from our network analyses were highly associated with immune-cell signaling and inflammatory/wound-healing pathways, especially tumor necrosis factor *α* (TNF*α*)-mediated nuclear factor *κ*B (NF-*κ*B) signaling (Xia et al., 2014; Taniguchi and Karin, 2018). Most of the functional associations we identified are involved in lipid metabolism, immune cell dynamics/homeostasis, cancer growth, cellular cargo trafficking, and the TNF*α*-NF-kB pathway (Figure 6C). A significant number of human cancers have constitutive NF-κB activity due to the inflammatory microenvironment and various oncogenic mutations. NF-κB activity not only promotes tumor cells proliferation, suppresses apoptosis, and attracts angiogenesis, but it also induces EMT plasticity which facilitates distant metastasis and adaptive cell state transitions (Xia et al., 2014; Taniguchi and Karin, 2018; Hutzen et al., 2019). Previous studies have also shown that the feedback system between TNF*α* and NF-*κ*B display intracellular oscillatory dynamics (Tian et al., 2015), which is relevant to our results as oscillations are precursors to the self-organization of complex attractor dynamics (Tiana and Jensen, 2013; Strogatz, 2015). Interestingly, mathematical and computational modelling studies paired with live-cell imaging have shown that intracellular chaotic dynamics can emerge in the protein oscillations of TNF*α* and NF-*κ*B (Jensen and Krishna, 2012; Heltberg et al., 2019). These studies suggest that the emergence of chaotic attractors in cancer cell signaling, and cancer-immune cell dynamics may be an indicator of its complex adaptive behaviors such as therapy resistance, metastasis, and resilience (Itik and Banks, 2010; Letellier et al., 2013; Khajanchi et al., 2018; Heltberg et al., 2019).

MYC gene targets had the highest enrichment score followed by EMT plasticity pathways, both of which are critical regulators of glioblastoma phenotypic plasticity, cancer stemness, and metastatic potential. Interestingly, oxidative phosphorylation has a higher enrichment score than glycolysis indicating a preferential selection of metabolic regulation. MTORC1, TNF*α* via NF-kB, and TGF*β*, critical factors regulating cancer growth, tumor microenvironment homeostasis, and cell fate plasticity were also found to be highly expressed with a roughly three-fold enrichment score. Many of these identified signaling pathways are also critical for neuronal-gliomal network regulation of glioma growth and progression (De Silva et al., 2023). Further, these signals are key immune cells-mediated inflammatory pathways required for regulating the CSC niche and conferring CSC states (Plaks et al., 2015; Batlle and Clevers, 2017; Gonzalez et al., 2018; Zhao et al., 2021). In the context of pediatric glioblastoma, Proneural-mesenchymal transition (PMT) or in general, the progenitor neuronal-glial transition at the developmental origin of pHGGs (Jessa et al., 2019), can be considered as the EMT analogues of phenotypic plasticity dynamics in between the heterogeneous cellular states. We predict the TPT1-PTPRZ1 feedback loop in coordination with central network markers such as PDGFRA, EGFR, OLIG1/2, are involved in these phenotypic transitions.

To support these speculations, follow-up studies using fractal dimension analysis of the reconstructed glioblastoma cell fate attractors in addition to attractor embedding from time-series single-cell datasets followed by chaos detection tools such as (positive) Lyapunov exponents and a positive metric/topological entropy should be applied to further test the presence of chaotic dynamics in glioblastoma cell fate decisions. We further speculate the upregulated pathways and signaling proteins identified by CellChat analysis in glioblastoma-immune dynamics may be indicative of potential transcription programs for reprogramming glioblastoma phenotypes towards stable neuronal lineages.

Graph-network algorithmic complexity was used as a cross-validation tool to identify causally related high centrality nodes and links in the transcriptional networks. Table 2 shows that the highest graph network complexity (*K*(*G*)) changes in the Bayesian networks was observed with the perturbation to MTRNR2L1/2 and OLIG1, which were identified as a feedback loop/circuit in our NLNET analysis (Figure 2E-F). Therefore, we found some equivalence between the *K*(*G*) of a Bayesian inference network and the NLNET algorithm. Further, the algorithmic complexity analysis on the Block Decomposition Method changes (i.e., K-complexity shifts) in the perturbation of LEAP networks (Table 3) agreed with the high information flow regulators/nodes identified using graph-theoretic network centrality measures. Therefore, our analyses suggest that algorithmic information dynamics and K-complexity is a robust measure of network dynamics steering glioblastoma cell fate dynamics/behaviors and can be used to predict causal relationships in the complex network patterns we inferred. S100B, a glial inflammatory and astrocyte marker and OLIG1/2 were identified as the top-ranking network markers with the greatest algorithmic complexity change (BDM difference) in the network links/edges’ perturbation analysis, in both network metrics (Bayesian and LEAP) as shown in tables 2 and 3, respectively. Thus, we conclude that OLIG1/2 and S100B are critical regulators of the causal relationships/algorithmic information dynamics of the inferred glioblastoma networks.

Together, our results demonstrate the use of complex systems approaches for understanding glioblastoma cell fate decisions and phenotypic transitions as complex adaptive processes. By studying the underlying attractor dynamics and regulatory networks driving these complex processes through our approaches, we were able to identify robust signaling pathways and transition genes acting as master regulators of the glioblastoma cell fate decisions. The complex systems methods deployed here suggest potential ways to reprogram CSC fates within aggressive tumors like glioblastoma towards benignity, which would have significant impacts on patient outcomes. This is further supported by the fact that the critical network markers (transition genes) we identified in our study are involved in glioma differentiation and cell fate commitments to neuronal and glial phenotypes (see Figure S8 in Supplementary Information). Further, our approach paves the way towards identifying clinically relevant targeted therapies against the adaptive, therapy resistant CSCs within glioblastoma, making it among the first in the emerging discipline of computational immuno-oncology and systems medicine.

Limitations of our study include the technical barriers of the scRNA-Seq data, i.e., batch effects, data sparsity, dropout events, and noise. The lack of immunophenotypic markers on T cells in the scRNA-Seq datasets further limited our analyses of immune-glioblastoma dynamics. Prospective studies should consider both immunophenotyping the infiltrated immune cells within glioblastoma tumors and longitudinal blood-immune profiling for liquid biopsy markers to better characterize the complexity of cancer-immune network dynamics. The absence of time-resolved data presents a greater barrier to the analysis of cell fate behavioral patterns as dynamical systems. While data constraints present a fundamental challenge in single-cell pattern discovery and data science, the complexity of analytical tools can also be limiting. For instance, the network inference methods employed here are restricted to statistical methods such as correlation metrics and Bayesian inference that may not be able to capture the complex dynamics of cellular differentiation networks when presented with time-resolved data. Although K-complexity estimates provided by the BDM-based network perturbation analysis provide a more rigorous network metric, they too were limited by binary thresholds and the binarization of network interactions. Going forward, K-complexity metrics should be expanded to fuzzy systems and continuous models of weighted networks. The lack of a rigorous network inference metric remains a central problem in causality inference. Furthermore, time-resolved datasets or experimental assays are required to distinguish positive and negative feedback loops in the inferred regulatory networks.

Our study was restricted to IDH-WT glioblastoma samples obtained from gene expression data from Neftel et al. (2019). Future studies should extend our computational methods to epigenetic sub-types of pediatric high-grade gliomas and histone variants of pediatric glioblastoma such as H3K27M and H3.3G34V/R to better elucidate how the identified feedback mechanisms may be involved in the broader context of glioma development and differentiation (Schwartzentruber et al., 2012; Boileau et al., 2019; Jessa et al., 2019). Prospective studies should also test other validation cohorts and compare the differentiation network dynamics amidst different pediatric glioblastoma subtypes including IDH-mutant (rare) samples, and oncohistone (epigenetic) variants such as H3.1K27M (prevalent in DIPG) and H3.3K27M with dysregulated polycomb repressor complex dynamics. The integration of multimodal profiling techniques (single-cell multiomics, proteomics, metabolomics, ChIP-Seq data, 3D-chromatin conformation capture by Hi-C mapping, single-cell chromatin modifications/nucleosome profiling, and methylome sequencing) may provide richer insights into the transcriptional patterns identified in glioblastoma cell fate differentiation dynamics and behavioral patterns. Our network analyses can be extended to multiscale networks including multiomic networks (i.e., protein-protein interaction networks, metabolic networks, epigenetic networks, etc.) to study the physiological/multicellular networks orchestrating collective cell fate decisions in glioblastoma plasticity dynamics.

Despite these limitations, our complex systems tools provided important insights into the central regulators of information dynamics in glioblastoma plasticity/differentiation networks. Importantly, the identified key transition genes showed overlapping results confirming the robustness and validity of our findings. Network centrality measures of the identified hub genes were similarly found to overlap with the K-complexity shift markers, which is indicative of causal relations in the glioblastoma networks. Nonetheless, though the robustness of the network transition markers we identified regulating glioblastoma differentiation dynamics across their transcriptional state-space (attractor landscape) depends on the hyperparameter tuning of the used lineage tracing/trajectory inference algorithms and the network inference metrics, the markers we identified in this study strongly overlap with the therapeutic targets/transition markers we identified in our previous network analyses using a different set of Waddington landscape reconstruction algorithms and network inference metrics on single-cell glioblastoma transcriptomic datasets (Uthamacumaran and Craig, 2022).

Our study underlines complex systems approaches as emerging tools for inferring causal relationships between the driver gene networks orchestrating adaptive cellular behaviors in glioblastoma ecosystems and can quantitatively characterize glioblastoma cell fate dynamics. Despite our use of different types of network inference algorithms, similar biomarkers were identified as critical regulators and drivers of glioblastoma transcriptional dynamics. As such, we can conclude that the inferred patterns of network dynamics and network biomarkers are robust transcriptomic signatures indicative of glioblastoma cell fate dynamics. The inferred glioblastoma networks are robust patterns classifying glioblastoma cell fate behavioral dynamics and may be clinically relevant targets for precision oncology.

## DATA AND CODES AVAILABILITY STATEMENTS

23,658 genes and 1943 cells (Neftel et al., 2019) GitHub page: Pending *

CALISTA: https://www.cabselab.com/calista

Hopland: https://github.com/NetLand-NTU/HopLand

MuTrans: https://github.com/cliffzhou92/MuTrans-release

Slingshot: https://bioconductor.org/packages/devel/bioc/vignettes/slingshot/inst/doc/vignette.html

CellChat: https://github.com/sqjin/CellChat

AR1MA1-VBEM: https://github.com/mscastillo/GRNVBEM

Julia LightGraphs: https://github.com/JuliaGraphs/LightGraphs.jl

CCAT (part of the SCENT-R package): https://github.com/aet21/SCENT

LEAP: https://cran.r-project.org/web/packages/LEAP/index.html

NLNET: https://cran.r-project.org/web/packages/nlnet/index.html

PHATE: https://github.com/KrishnaswamyLab/PHATE

Community Structure Detection: https://www.r-bloggers.com/2020/03/community-detection-with-louvain-and-infomap/

ReactomeGSA: https://bioconductor.org/packages/release/bioc/vignettes/ReactomeGSA/inst/doc/analysing-scRNAseq.html

Molecular Signature Databases GSEA: https://crazyhottommy.github.io/scRNA-seq-workshop-Fall-2019/scRNAseq_workshop_3.html

https://stephenturner.github.io/deseq-to-fgsea/

OACC (for Network Complexity Perturbation Analysis): http://complexitycalculator.com/

https://github.com/algorithmicnaturelab/OACC (Zenil et al., 2018)

