## Supplemental Info for "Cell fate dynamics reconstruction identifies TPT1 and PTPRZ1 feedback loops as master regulators of differentiation in pediatric glioblastoma-immune cell networks"

**Abicumaran Uthamacumaran**

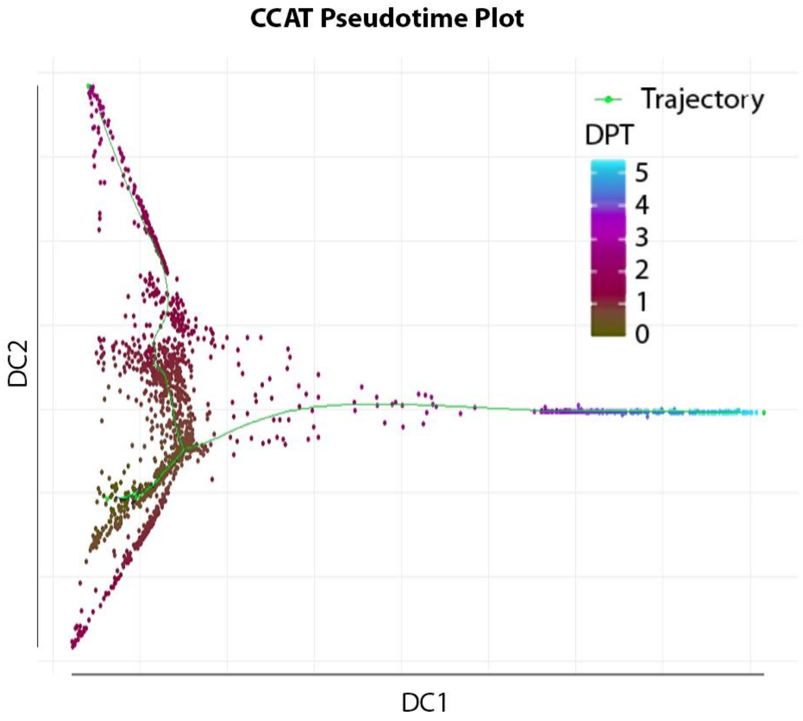


**Figure S1. CCAT identifies bifurcation in glioblastoma cell fate differentiation dynamics.** The bifurcation of glioblastoma cell fate lineages from inferred root cells are displayed by the CCAT algorithm in diffusion map space. The cell fates in the CCAT mapping are colored by their diffusion pseudotime (DPT) inferring the lineage trajectories of these cell fate transitions, where blue indicates high DPT potential (i.e., higher differentiation capacity) and red/green indicates low DPT potential. The CCAT algorithm was used as a validation tool to verify the cell fate attractors observed in Figure 2. Two main cell fate trajectories are identified starting at the identified root-cells with the highest entropy score (stem cell-like states), with each trajectory characterizing a differentiation pathway to committed phenotypes.

**PHATE algorithm maps glioblastoma cell fate differentiation by marker expression of critical transition genes.**

Glioblastoma cell fate differentiation was mapped using PHATE (Moon et al., 2019) from the critical cell fate regulators identified from our network analyses (see description of the algorithm below). As seen in Figure S2, the progressive clustering we found is indicative of differentiation, which usually does not result in distinct clusters. A clear progression from right to left in PHATE1 seems to correlate with a differentiation progression. We found that PABPC1 (Figure S2A), MTRNR2L1/2 (Figure S2C and S2D), PSAP

(Figure S2H), and TPT1 (Figure S2E) displayed increasing expression as the glioblastoma cell fates transition along PHATE1 dimension. TPT1 and MTRNR2L1/2 had the greatest and smoothest marker expression along the transition dynamics. These findings agree with the interactions identified in our complex network analyses. The expression patterns of OLIG1 (Figure S2B), S100B (Figure S2F), and PTPRZ1 (Figure S2G), determined to be critical regulators of glioblastoma-immune transcriptional networks through network measures (BDM scores) were found to be similar as they all lack high-expression of the gene biomarker within the cell cluster on the higher positive end of the PHATE1 (horizontal) axis.

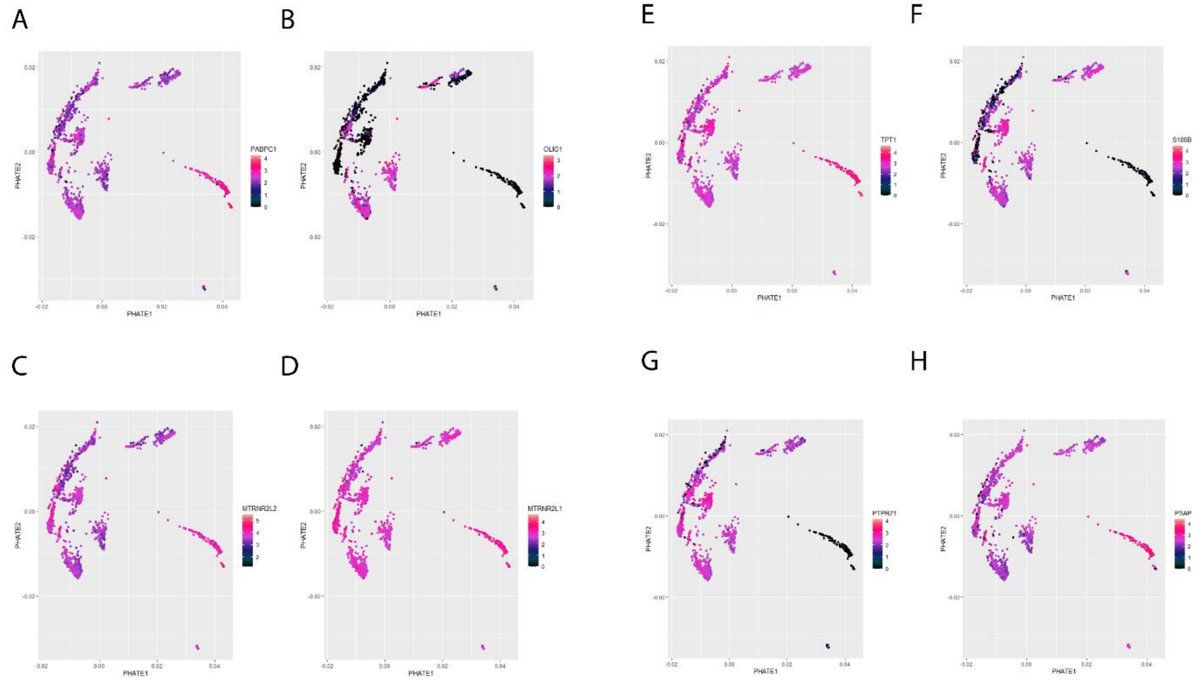


**Figure S2. PHATE cell fate trajectory analysis of glioblastoma cells.** The differentiation map of glioblastoma cells plotted for selected marker genes: A) PABPC1, B) OLIG1, C) MTRNR2L2, D) MTRNR2L1, E) TPT1, F) S100B, G) PTPRZ1, and H) PSAP. The expression scale bar indicates null expression (dark indigo) to maximal normalized expression of 4 (pink). Violet indicates a medial expression between the two.

**Network medicine and trajectory inference algorithms identify causal patterns in the complex signaling dynamics orchestrating glioblastoma cell fate decision-making**

We reconstructed the minimum spanning tree (MST) of the Bayesian (Figure S3A) and LEAP (Figure S3B) glioblastoma networks using the Kruskal algorithm in the LightGraphs Julia package (see Methods in the Main Text). The MST provides a network topography optimized for the minimum path between the nodes by their respective weights (i.e., signaling strength). In the Bayesian network, the heaviest weight edge (interaction strength of 1.0) was found between MTRNR2L2 and PTPRZ1 followed by the interaction between TPT1 and MTRNR2L1 with a weight of -0.642, indicative of a negative feedback system (Fig S3A). Other strong positive feedback interactions identified were between SOX6 and MTSS1 (with a weight of 0.186) and that between GATA2 and EMP1 (weight of 0.133). These relationships were also identified as bridges of the network. MTRNR2L2 was identified as the center of the network. These findings, cross validated by Prim's algorithm, are in good agreement with the nodes with the highest network centrality measures identified in our results. The MST of the LEAP network showed the highest edge weight of 0.896

for the interaction between OLIG2 and EGFR, followed by an interaction weight of 0.818 between SOX6 and PTPRZ1 (Fig S2B). EGFR was found to be the center of the network.

We performed two types of dimensionality reduction with respect to the glioblastoma cell lineage trajectories: diffusion component map (DC) and principal component analysis (PCA) (Figure S3 C-D) and found a causal structure with a bifurcation in both pattern spaces. As seen in Figure S3C, glioblastoma cells were found to branch into two cell fate commitments in the diffusion map space with a smoother bifurcation observed in the PCA space (Fig S3D). This bifurcation confirms the inferred complex attractor observed in the Waddington landscape reconstruction algorithms mapping glioblastoma cell fate dynamics (Figure 2 in the Main Text). The results shown in Figure S3E-F further support an underlying causal order (or complex attractor) steering glioblastoma cell fate bifurcations. This attractor closely matches the CALISTA attractor and the CCAT-colored cell fate trajectory shown in Figure S1. Hence, our findings collectively suggest the presence of a complex attractor, with a pitchfork bifurcation steering glioblastoma cell fate decisions and phenotypic transitions/plasticity.

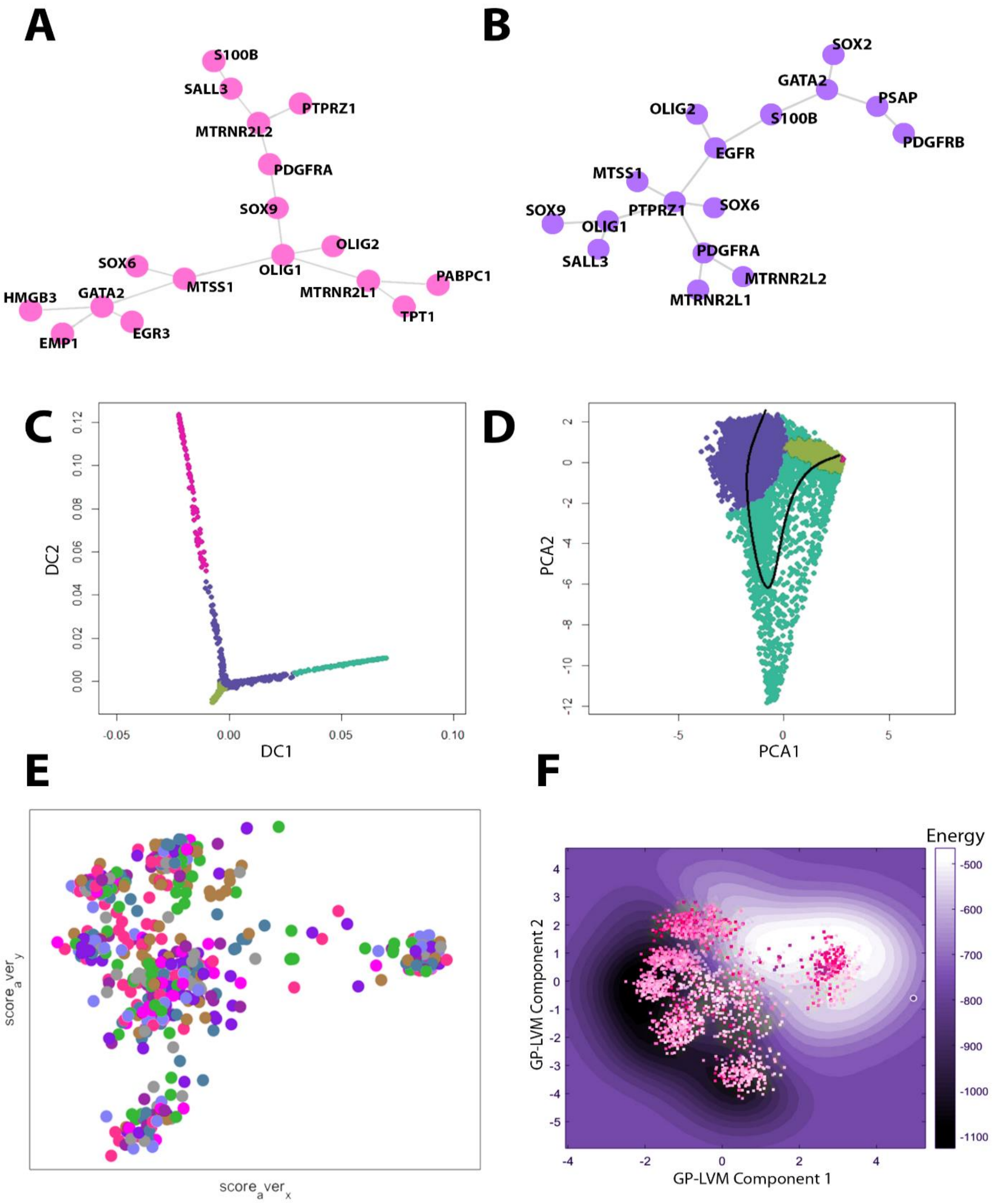


**Figure S3. Algorithmic reconstruction of glioblastoma cell fate trajectories and decision-making.** A) Minimum Spanning Tree (MST) of the Bayesian glioblastoma network performed by the Kruskal algorithm. B) MST of the LEAP glioblastoma network by the Kruskal algorithm. C) Diffusion map of the glioblastoma cell fate lineages mapped by Slingshot algorithm. D) glioblastoma cell fate trajectory mapped by Slingshot algorithm in PCA dimensionality reduced space. The colors indicate distinct heterogeneous phenotypes identified by Slingshot within the glioblastoma population. The black curve corresponds to the differentiation map of cell fates amidst the distinct phenotypes. E) Two-dimensional contour plot of the glioblastoma cell fates on the MuTrans attractor landscape. F) The contour plot of the reconstructed Waddington's epigenetic landscape from the Hopland algorithm in GPLVM dimensionality reduced space. The energy level of the landscape topography is shown by the color gradient bar from black indicating low (negative) energy to white indicating high (positive) energy. The glioblastoma cells are colored in a gradient from pink to white, where pink denotes high expression of critical transition genes identified in our network analyses (LEAP and Bayesian networks in Figure 4A-B in the Main Text) and white denotes low expression.

**Nestedness and distinct modules are present in glioblastoma gene regulatory networks**

Community detection algorithms from the igraph R-package (Csardi and Nepusz, 2006) revealed modularity within the complex GRNs. Similar clusters of genes were seen in the three distinct modules in both the Bayesian and LEAP networks (Figure S4). The glioblastoma stemness markers identified by Suva et al. (2014), including the transcription factors SALL3, OLIG2, and SOX, were found to be clustered into a single module in both networks. Whether the three distinct modules are related to the transcription programs conserved within the glioblastoma trilineage reported by Couturier et al. (2020) remains unclear.

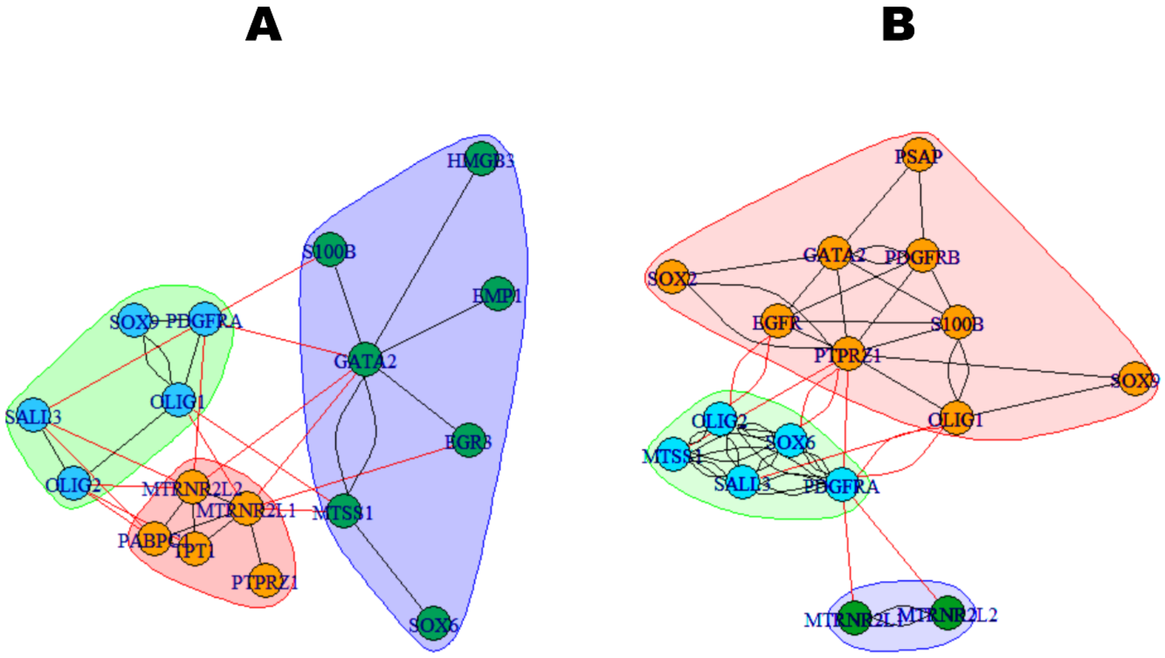


**Figure S4. Community structure detection on glioblastoma networks.** The Louvain community structure detection algorithm were assessed as modularity optimization on A) Bayesian Network, and B) LEAP Network. The colored maps indicate distinct communities (modules) within the complex networks. Identical community clusters were also obtained for the Infomap algorithm.

**CD4+ subset-specific differential markers are discovered in the PCA clustering of glioblastoma-infiltrated T cell population**.

Since the glioblastoma-infiltrated T cells were not immunophenotyped, we pooled the T cell population and performed clustering expression analysis using the Seurat algorithm. As most immune-specific cell surface markers/receptors must be transcribed, we interpreted mRNA transcript levels detected in the scRNA-Seq counts to be indicative of their relative protein abundance/expression. As shown in Figure S5, the PCA loadings of the normalized counts revealed markers like GRN and CX3CR1 which were observed in the CellChat signalling inference analysis. Further, MCH-class II histocompatibility complex-associated proteins like HLA-DRB1 and HLA-DRA were highly expressed in all T cells. PSAP, one of the differential markers identified in our single-cell analyses, was similarly highly expressed.

We found CD4 to be highly expressed in the T cells whereas there was a complete absence of CD8 (Figure S5). The high expression of CD74 further confirms these are CD4+ T cells, since CD74 is an HLA class II histocompatibility antigen required for CD4+ T cell activation and responses (Germain, 2002; Yakimchuk, 2016). CD74 is also a cell surface receptor for the cytokine macrophage migration inhibitory factor (MIF) that was identified in our CellChat analysis. The observed expression of CD14 is also a recognition pattern for the microglia/macrophages, suggesting cross talk and complex dynamics between tumor-infiltrated T cells and macrophages within the glioblastoma ecosystem. As seen in Fig S5B, the high expression of STAT3 and STAT6 indicates that we cannot infer the differentiated subsets of CD4+ T cells (Yakimchuk, 2016), and subset-specific interleukins did not show variable expression. However, we saw high TGF-β and STAT5-IL-2 signaling in the GSEA analyses (Figure 6 in the Main Text), both of which are critical for the differentiation of regulatory T cells (Tregs) (Mahmud et al., 2013) that are critical for immunosuppression and tumor immune escape/evasion in tumor ecosystems (Verma et al., 2019). Further classification of other GBM-infiltrated immune cell subsets such as the macrophages/microglia should also be investigated in prospective single-cell analyses.

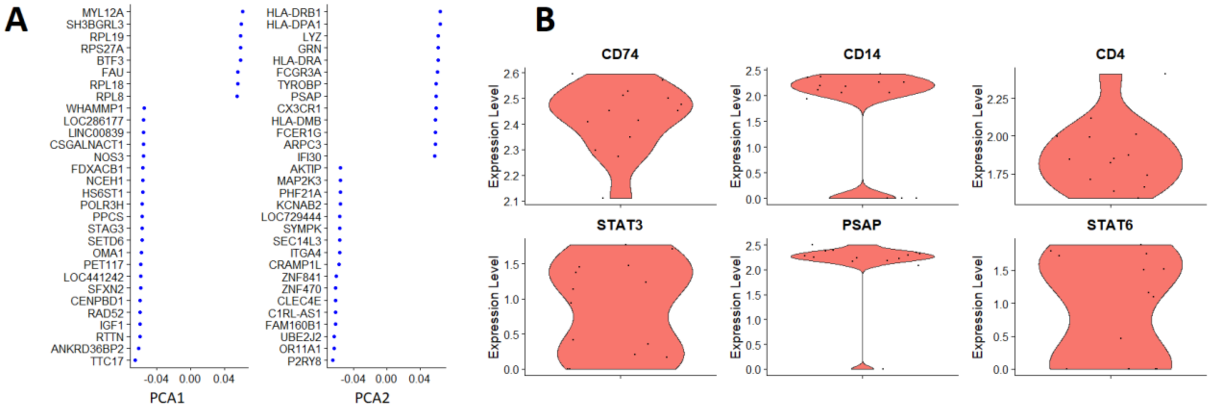


**Figure S5. T cell PCA analysis.** A) Differentially expressed markers in glioblastoma-infiltrated T cells in the PCA pattern space. B) Relative expression of T cell subset-specific markers in the PCA space. The width indicates the variabikity (standard deviation) and the height represents the expression level.

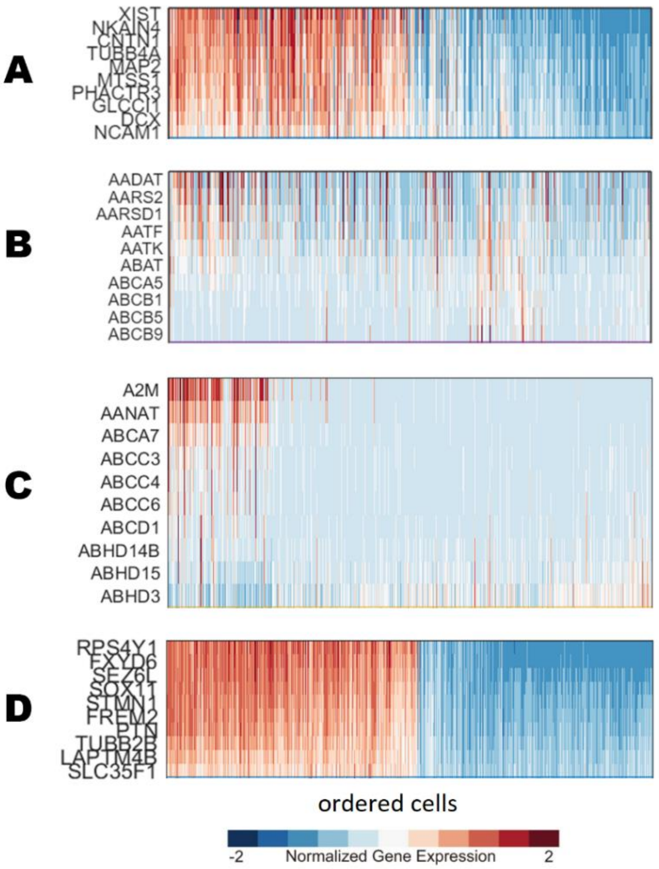


**Figure S6. MuTrans Heatmaps with top 10 transition genes (k=4).** A) Transition markers for cell fate transition from cluster 1 to cluster 2 on the landscape shown in Figure 5. B) Transition from cluster 1 to cluster 3. C) Transition from cluster 4 to cluster 1. D) Transition from cluster 2 to cluster 3. The transition markers in B and C were determined to be not as strong in expression were therefore omitted from the network analysis shown in Figure 5D to avoid false discovery.

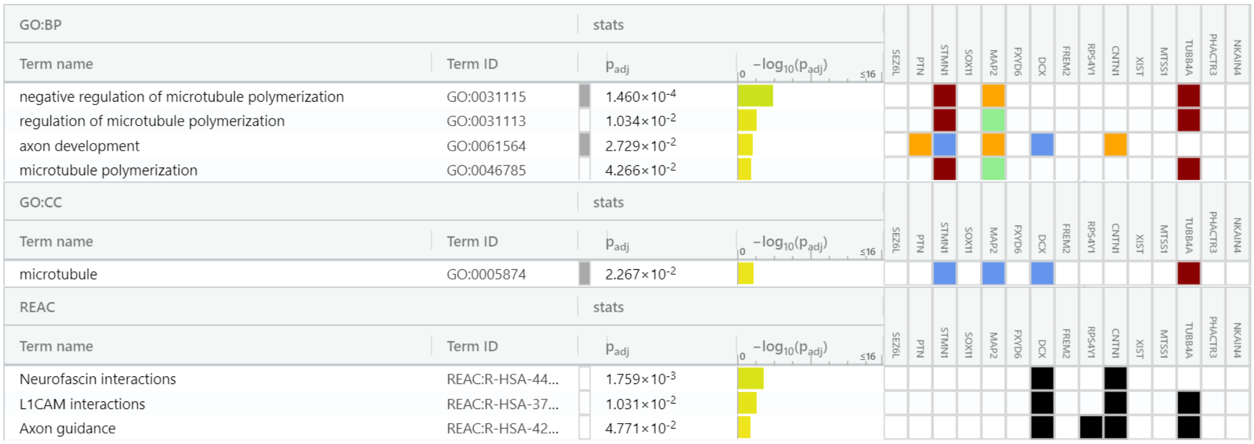


**Figure S7. gProfiler analysis of network markers from MuTrans algorithm**. A gene set enrichment of only the MuTrans network markers reveals they are involved in cytoskeletal reorganizations such as microtubule polymerization, and axon guidance, suggesting their involvement in phenotypic plasticity, glioma migration, and neuronal-glioma interactions.

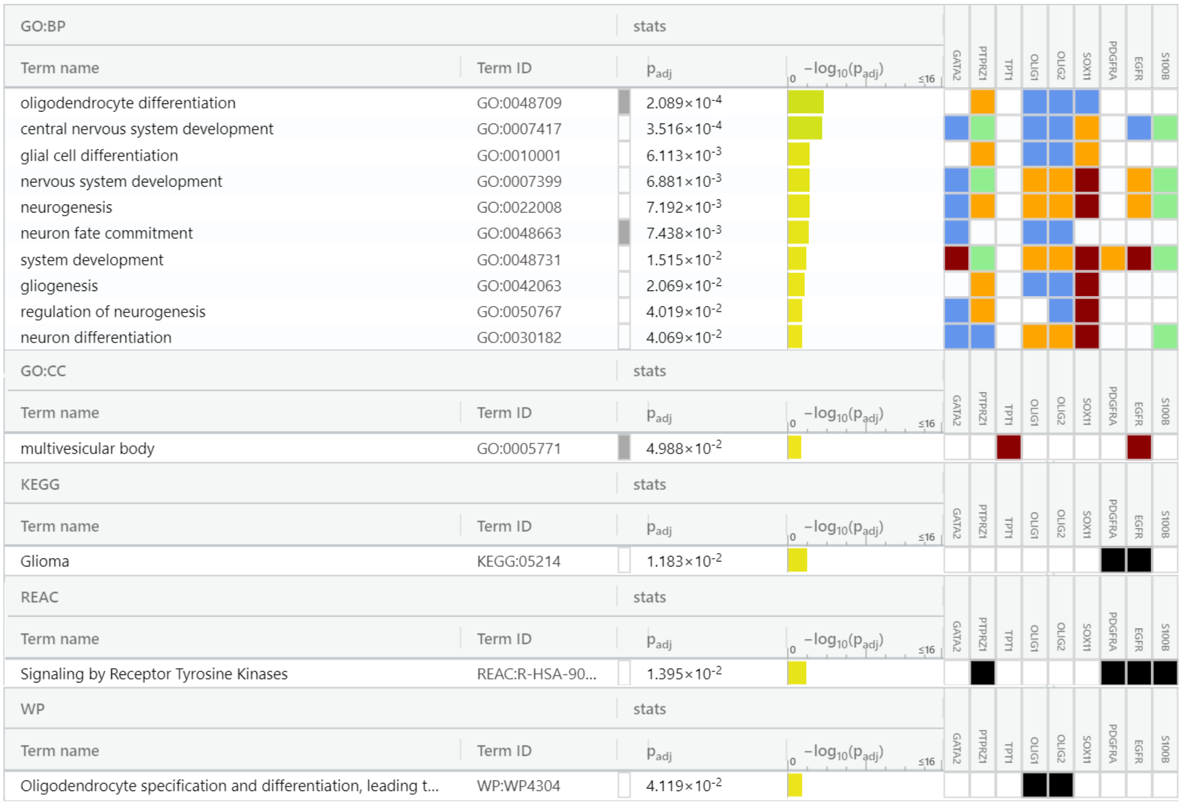


**Figure S8. gProfiler analysis for top network markers (transition genes).** The gene set enrichment analysis using gProfiler identified the critical regulators of our analyses to be involved in glioma differentiation and cell fate commitment dynamics. As seen, the transition genes we identified in our study are involved in various cell fate transitions such as oligodendrocyte differentiation, glial cell commitments, and neuronal fate commitments.

### ADDITIONAL DETAILS OF METHODS AND ALGORITHMS

#### Complex attractor dynamics

The cell fate patterns in transcriptional (gene-expression) state-space were shown to form complex attractors in our algorithmic analyses. Complex interaction networks of genes and proteins steering cell fate dynamics can exhibit temporal oscillations (Tiana and Jensen, 2013). A time-delay by, for instance, the formation of a protein complex or protein synthesis-degradation dynamics, can give rise to these temporal *oscillations*, examples of which include the expression of tumour-suppressor protein p53, NF-kB and Wnt, involved in cancer processes such as cancer (stem) cell fate decisions, regulation, and differentiation dynamics (Geva-Zatorsky et al., 2006; Jensen et al., 2010; Tiana and Jensen, 2013). Signaling oscillations correspond to the presence of more complex attractors in cancer signaling state-space, as indicated by the presence of bifurcation(s) (Prigogine, 1980; Janson, 2012; Strogatz, 2015). Complex attractors underlying collective cell fate choices in signaling/gene expression state-space may result in apparently random, irregular, or unpredictable behavioral patterns making the complex system difficult to treat (Strogatz, 2015). Previous studies have shown that cancer cell fates correspond to nonequilibrium, unstable attractors in cancer signaling state-space (Huang 2006; Huang et al., 2009; Fang et al., 2019). However, whether they are complex attractors exhibiting chaotic dynamics remains to be verified due to a general lack of appropriate time-series cancer datasets.

#### Clustering methods and trajectory inference algorithms

Multiple data science algorithms and statistical machine learning pipelines were used in this study, all consisting of a similar workflow for inferring cell fate dynamics from the scRNA-Seq datasets: 1) single-cell clustering based on some distance metric or correlation metric to infer a neighborhood graph between cells or genes, 2) differential gene expression analysis by use of pair-wise correlation metrics or similarity index, and 3) cell lineage reconstruction, and pseudotemporal ordering on a dimensionality reduced space (Nguyen et al., 2020). Some algorithms can further determine the transition genes steering the cell fate trajectories. Pseudotime is a quantitative measure of the flow of time in biological process (such as cell fate differentiation or phenotypic transitions) along which the cell fate dynamics are arranged based on the assumption that cells with similar gene expression profiles cluster together (Nguyen et al., 2020; Hu et al., 2020).

#### Seurat clustering

Seurat clustering was performed by embedding the cells with similar gene expression patterns onto a graph structure (e.g., the K-nearest neighbourhood (KNN) graph) based on the Euclidean distance in principal component analysis (PCA) dimensionality-reduction space (Stuart et al., 2019). The Jaccard similarity index was used to define the overlaps in local neighbourhoods of cell clusters (which may correspond to basins of attraction). Lastly, the graph was partitioned into highly interconnected clusters by use of iterative modularity optimization techniques such as the Louvain community detection algorithm or the smart local moving (SLM) algorithm (Stuart et al., 2019). Additional downstream analyses were performed on the modularity optimized (nested) cell clusters to identify key expression markers distinguishing them. We pooled the topmost differentially expressed markers for our network analyses. Further details of the parameters and clustering protocol are found in Uthamacumaran and Craig (2022).

**CALISTA**

Waddington landscape reconstruction measures the maximum likelihood of cells occupying a steady-state probability distribution of mRNA defined according to the stochastic two-state model of gene transcriptional process (Gao et al., 2020). For the reconstruction in CALISTA, the cell fate positions were determined by an iterative maximum likelihood clustering algorithm, used to generate a consensus matrix, followed by k-medoids clustering. A lineage progression graph was then constructed using cluster distances assessing similarity in gene expression amidst distinct clusters and linear interpolation. Lastly, transition genes between any two connected clusters in the cell lineage graph were extracted. These transition genes may correspond to candidate biomarkers regulating complex cell fate dynamics such as cell state transition during differentiation or cell fate interconversions.

**Hopland algorithm**

The Hopland algorithm first normalizes gene expression data and, by filtering out genes with low variances, selects differentially expressed genes to map cells onto a nearest-neighborhood graph reconstructed by Isomap, a type of manifold learning algorithm/nonlinear dimensionality reduction. The data is then analyzed using the Continuous Hopfield Network to reverse-engineer the topography of the Waddington's epigenetic landscape determined by the Lyapunov energy function. The probabilistic nonlinear dimensionality reduction Gaussian process latent variable model (GP-LVM) generates a mapping between the original high-dimensional data space and a 2D latent space (Guo and Zheng, 2017). Cell states with specific gene expression patterns were assumed to be fixed-point attractors, the simplest (most stable) of attractors in dynamical systems theory, and their time-evolution was mapped as a state-transition on the landscape. The geodesic distances between the cells were calculated by the fast-marching algorithm to estimate the pseudotime trajectory of the cell fate dynamics. Using the extracted geodesic distances as the weights of edges connecting the cells, a minimum spanning tree (MST) was constructed to demonstrate the cell fate bifurcations towards distinct attractors of the landscape. The CHN consists of N interconnected neurons representing the N genes.

The rugged landscape represents the Waddington's epigenetic landscape, wherein the differentiation processes of cells follow the trajectories of potential energy minimization resulting in the topography of the landscape. The transition cell fates are observed as flow trajectories between these basins of attraction. The dynamics of CHN system can be described by the Lyapunov energy function given by

$$E = -\frac{1}{2}\sum_{i=1}^{N}\sum_{j=1}^{N} W_{ij}U_iU_j + \sum_{i=1}^{N} I_iU_i + \sum_{i=1}^{N} \delta_i \int_0^{U_i} g_i^{-1}(u)\, du.$$

which is iteratively minimized as the CHN learns to recognize patterns from the gene expression dataset and store them. In the equation above, the gene expression of a cell is characterized by the outputs $V$ of the number of neurons (genes) $N$, $W_{ij}$ is the weight matrix representing the interconnection weight coefficient from neuron $j$ to neuron $i$, the external input $I_i$ represents a combination of propagation delays (noise in transcriptional regulation and other genes), $\delta_i$ denotes the degradation rate of the $i$-th gene, $g$ is the activation function (sigmoid function), $U$ are the energies of the neurons (genes). For a gene j, this energy is expressed by the sigmoidal function

$$U_j = g_j(V_j) = \frac{1}{\left(1 + e^{-\frac{V_j - \mu_j}{\sigma_j}}\right)},$$

where $\mu_j$ and $\sigma_j$ are the mean and standard deviation of the expression levels of the $j$-th gene in all cells, respectively (Guo and Zheng, 2017).

**MuTrans algorithm**

MuTrans uses coarse-grained transition rate theory and Langevin dynamics to map the complex bifurcation dynamics of cell fate decisions and capture intermediate, hybrid (mixed) cell states during cell state transitions. The transition trajectories can distinguish meta-stable and transition cells by assigning them to basins of attraction on the dynamical manifold (Waddington/attractor landscape). MuTrans assumes the cell-fate dynamics and phenotypic interconversions can be modelled by random walks (Brownian motion) among individual cells through the random-walk transition probability matrix (rwTPM) (Zhou et al., 2021). The cell-to-cell rwTPM was generated from coarse-grained dynamics, by assigning cell distributions and modeling the transitions as the Markov Chain among clusters with the transition probability matrix (created attractor basins). This process resulted in an optimization problem of cluster-cluster and cell-cluster assignment. Cell lineages and transition trajectories were inferred from the most probable path tree (MPPT) approach or maximum probability flow tree (MPFT) approach. The distance between cell clusters was then optimized by the quasi-Newton method. Both, Hopland and MuTrans are Gaussian mixture models (GMM)-based attractor landscape reconstruction algorithms in tracing cell fate dynamics.

To visualize cell fate decisions, MuTrans reconstructs an attractor landscape by soft clustering in low dimensional space towards basins of attraction. Metastable cell states are assumed to cluster around fixed-point stable attractors. The Fokker-Planck equation of the over-damped Langevin equation (OLE) is used to map the cell fate dynamics as the time-evolution of a probability distribution and determine the steady-state solutions near stable fixed-points (attractors). A global attractor landscape was obtained by fitting a Gaussian Mixture Model (GMM) constructing a two-dimensional Gaussian probability distribution density P(z), to obtain stationary distribution of coarse-grained cell fate dynamics. Hence, with the assumption that the single-cell data is obtained from a probability distribution $\rho(x)$ with a Boltzmann-Gibbs density, i.e., $\rho(x) \propto e^{-\frac{U(x)}{\varepsilon}}$, the cell fate dynamics and transitions/interconversions were modelled as microscopic random walks by approximating the OLE given by: $dX_t = -\nabla U(X_t)dt + \sqrt{2\varepsilon}dW_t$ , where $dX_t$ is the differential of the position-vector state X (of the cell) with respect to time, $\varepsilon = k_B T$ (thermal noise term), W denotes the Wiener process, and $\Delta U$ denotes the barrier height of the transitions (energy difference between the attractor and its saddle point) (Zhou et al., 2021). In the small noise regime, the OLE dynamics are reduced to Kramer's rate formula in transition state theory. The energy of individual cell fates on the attractor landscape was given by the potential energy $U = -\ln P(z)$.

**Entropy landscape inference by CCAT**

Correlation of connectome and transcriptome (CCAT), part of the SCENT-R package, is a single-cell potency measure that can estimate the normalized signaling entropy rate (SR) of cell populations (Teschendorff and Enver, 2017; Teschendorff et al., 2021). We used CCAT to infer the dynamic lineage trajectories and cell state transitions on a diffusion map space. The entropy rate is a measure of the cells' differentiation potency. CCAT thus defines the transition probability between cell fate clusters based on their entropy difference. The log-normalized scRNA-Seq data was first filtered to avoid zero values in the data matrix, and the normalization accounted for a pseudocount of 1.1, so that the log (counts + 1.1) transformation takes on a minimum value above zero. The CCAT scores were assessed on the transformed expression matrix. The root-state was inferred from the top 97 cells with highest potency using the *InferDMAPandRoot* function (a destiny package in R to construct the diffusion map and Markov Chain transition matrix over all cells). The diffusion map was constructed from 5211 variable genes. The transition probabilities were calculated using the transition matrix over a kNN graph, followed by eigengap decomposition and a running walk-trap community detection algorithm which identified a root-state consisting of 31 cells. The cell transitions/trajectories on the diffusion map were colored by CCAT (Figure S7).

Given the log-normalized scRNA-Seq profile (transcriptome) $x = \{x_1, \ldots, x_G\}$, where $G$ is the number of genes in a regulatory graph network, let $A$ be the corresponding $G\ X\ G$ adjacency matrix (Teschendorff and Enver, 2017). Then, we can define a stochastic diffusion matrix P on this graph network, by the entries: $p_{ij} = \frac{A_{ij}x_j}{\sum_k A_{ik}x_k}$, where $A_{ii} = 0$ and $A_{ij} = 0$ if genes $i$ and $j$ are not neighbors in the network, and $A_{ij} = 1$ if they are.

The entropy rate (SR) is then defined as $SR = -\sum_{i,j} \pi_i p_{ij} log p_{ij}$, where $\pi$ is an invariant measure of the Markov Chain process on the graph satisfying $\pi P = \pi$. By taking a global mean-field approximation, we defined a parameter $k$ as the connectome that describes the degree/connectivity of the node(gene) $i$ in the network (Teschendorff and Enver, 2017). Then, CCAT is defined by the Pearson Correlation Coefficient (PCC) relating the transcriptome x and the connectome k, given as $CCAT = PCC(x, k)$. Since CCAT is a PCC between the connectome and transcriptome of a cell, it can take on values between -1 and 1, with increasing values indicating higher differentiation potency.

**Slingshot**

Slingshot is a pseudotemporal reconstruction and cell lineage inference algorithm (Street et al., 2018) that, given a gene expression matrix, assumes that the $N$ cells have been partitioned into $K$ clusters, potentially mapping distinct cellular states. We then chose a dimensionality reduction step (e.g., diffusion map or principal component analysis (PCA)). Pseudotemporal reconstruction was performed by curve-fitting using Euclidean (or related) distances on the clustered cells. Slingshot identifies cell lineages by treating cell clusters as nodes in a graph and tracing a minimum spanning tree (MST) between the nodes. Constructing an MST involved a distinct distance measure between nodes (cell clusters) known as the Mahalanobis-like distance, i.e., a covariance scaled Euclidean distance, which accounts for the cell cluster shapes (Street et al., 2018).

**PHATE clustering for cell fate transitions mapping**

Potential of heat diffusion for affinity-based transition embedding (PHATE) is a tool for visualizing a denoised map of phenotypic transitions and differentiation (cell fate progression) in high dimensional single-cell data (Moon et al., 2019). Given an input gene expression matrix, PHATE 1) computes the pairwise distances from the matrix, 2) transforms the distances to affinities, 3) learns global relationships via the diffusion process, and 4) uses a potential distance to encode these learnt transition maps into a low-dimensional embedding. The embedding allows for the inference of local and global dynamic structures in the gene expression patterns (Moon et al., 2019).

In the first step, PHATE uses a Gaussian kernel-manifold learning on all pairs of points to transform the global Euclidean distances into local affinities to identify similarity in cellular gene expression profiles. Then a Markovian random-walk diffusion process is used to construct the global transition structure (Moon et al., 2019). The diffusion time scale parameter $t$ determines the number of steps taken in the random walk embedding and is optimized using von Neumann entropy. In our analyses, $t$ was set to its default value. Local branching points were determined in cell fate progression/differentiation process by use of a k-nearest neighbor (kNN) graphing algorithm. Lastly, EMD score analysis provided differential expression analysis to identify robust biomarkers in the cell fate switches.

**NETWORK MEDICINE ALGORITHMS**

**Bayesian network inference (AR1MA1-VBEM)**

A Bayesian network is a graph-theoretic model that represents probabilistic relationships (the edges of the network) between differential gene markers (nodes) identified from clustering algorithms. The network topology can be described by a set of binary variables, where $x_i(j) = 1$ specifies that the $j$-th gene is a parent of the $i$-th one (whereas $x_i(j) = 0$, otherwise). The activation or inhibition regulatory interactions amongst the genes are also inferred by the algorithm. Given a set of gene expression values from a gene expression matrix, the AR1MA1 algorithm estimates the most probable value of a posteriori of a set of parameters and hidden variables describing the network interaction dynamics (i.e., identifying relationships between a pairs of genes) that fits the Bayesian likelihood function given by Bayes' theorem (Sanchez-Castillo et al., 2018). Noise in the gene expression is modelled by a multivariate Gaussian with unknown mean and variance. Since the computation of the posterior probability of interactions from the likelihood function can be intractable for complex datasets (i.e., NP-hard problem), the VBEM method optimizes a functional that depends on a free distribution of the hidden variables and parameters describing the system (i.e., the gene expression counts). By iterating the VBEM method, the algorithm converged towards a network topology characterizing the probabilistic relationships expected in the gene regulatory network. The weights (edges) of the network were computed as the product of the posterior probability and the weight of each probability (normalized to the maximum value).

**Modularity detection by NLNET**

NLNET is a modular network inference algorithm consisting of four methods: DCOL-based K- clustering, non-linear network reconstruction, non-linear hierarchical clustering, and variable selection for generalized additive model. First, given a gene expression matrix, NLNET computes the pairwise relations between genes using DCOL. Next, gene-wise false discovery rate inference is performed to determine significant edges (links) for each gene. DCOL obeys a normal distribution under the null hypothesis that two genes are independent of one another (Liu et al., 2016). A gene-specific null distribution is obtained through random permutations (n=500 times). The gene distance matrix is determined using DCOL and p-values are calculated for tests of the difference between gene relationships and the normal distribution. We processed the vector of p-values using the *fdrtool* package to obtain local false discovery rate values. Dynamic thresholding of these false discovery rate values allows for the reverse-engineering of the network topology (connections between the genes). The community detection via multi-level optimization of modularity and label propagation was used to decompose the inferred gene network into communities by modularity optimization (Liu et al., 2016).

**Graph network complexity**

K-complexity is a robust measure of a network's complexity (Zenil et al., 2016). K-complexity ($K(G)$), also known as Kolmogorov or algorithmic complexity, quantifies the shortest length of a computer program (in bits) required to describe a graph network ($G$ ). $K(G)$ is analogous to Shannon entropy as a measure of complexity (i.e., a lack of randomness) (Zenil et al., 2019) but it is a more robust tool than Shannon entropy to measure the complex dynamics of networks. Although Shannon entropy can quantify the amount of information in a complex system or network, it is not informative on causal connections nor does it provide any insight into the algorithmic content of a graph network.

Formally, the Kolmogorov complexity of a discrete dynamical system for a string, array, or graph/matrix, $G$ is given by

$$K(G|e) = \min\{|p|: U(p,e) = G\},$$

where $p$ is the program that produces s and halts running on a universal Turing machine $U$ with input $e$. $K(G)$ is the length of the shortest description of the generating mechanism of the network or system. By the invariance theorem, $K(G)$ is a function that takes a graph network or matrix $s$ to be the length of the shortest program $p$ that generates $s$. A graph network or system is defined as random (non-causal) if $K(G)$ is about the same length in bits of $G$ itself. A network with low Kolmogorov complexity is then defined as c-compressible if $|p| + c = |G|$ and the graph G is random if $K(G) \approx |G|$.

$K(G)$ is incomputable and must be approximated using tools from algorithmic information dynamics, the most robust estimate of which is obtained by the block decomposition method (BDM). An R implementation of the BDM from the online algorithmic complexity calculator (OACC) and its GitHub page are provided in the Code and Data Section. BDM is defined as

$$BDM = \sum_{i=1}^{n} K(block_i) + log_2(|block_i|),$$

where the block size must be specified for the n-number of blocks. When block sizes are higher, better approximations of the K-complexity are obtained. For smaller block sizes, BDM works analogous to Shannon entropy (Zenil et al., 2019). Here, we used algorithmic perturbation analysis on the Boolean glioblastoma networks to determine which nodes and links are central regulators of the complex networks by their shift difference in graph network complexity.

**CellChat**

CellChat infers signaling networks through network measures and centralities from graph theory and manifold learning. First, differentially expressed signaling genes within an scRNA-seq dataset are determined using the Wilcoxon rank sum test with a significance level of 0.05. Next, the ensemble average cell expressions is computed (Jin et al., 2021). To calculate the intercellular communication probabilities, CellChat models the ligand-receptor mediated signaling interactions using the law of mass action and assumes these obey a random walk-based network propagation, with a Hill function modelling the interaction kinetics with a parameter $Kh$ (whose default value was set here to be 0.5). Statistically significant interactions between distinct cell groups are identified using a random permutation test.

To identify key communication patterns within the signaling networks, CellChat uses an unsupervised learning method non-negative matrix factorization (Jin et al., 2021). Similarity amongst the intercellular communication network patterns is assessed using the Jaccard similarity measure and the calculation of a functional similarity matrix S. Manifold learning on the signaling networks is then performed in three steps (Jin et al., 2021). First, a shared nearest neighbor (SNN) similarity network of the signaling pathways is constructed by k-nearest neighbors. Second, cells are classified into distinct groups, and the SNN graph of cells is embedded onto a low dimensional pattern space by principal component analysis (or diffusion map). Lastly, cells are clustered by modularity optimization of the SNN graph through the Louvain community structure detection algorithm. The cell density of each group is determined by an eigenvalue decomposition spectrum by analyzing the Laplacian matrix derived from the Louvain algorithm.

Graph-theoretic centrality measures in weighted-directed networks, including outdegree, in-degree, flow betweenness and information centrality, were used to identify key senders, receivers, mediators, and influencers for the intercellular signaling communications (Jin et al., 2021). For weighted-directed networks, weights were computed as communication probabilities using the gene expression counts. The flow betweenness measures the cells' tendency to control information/communication flow in the signaling networks, while the information centrality score was a hybrid of the closeness and eigenvector centralities of signaling networks, wherein a higher value indicates a greater control on the information flow (Landherr et al., 2010; Jin et al., 2021).

**Network centrality measures**

Centrality measures assess the degree of influence graph-theoretic network nodes or structures have on the complex network. As such, these measures inform on the critical nodes governing the information dynamics across a complex network and may provide robust markers, features, or patterns for identifying and controlling/reprogramming disease network dynamics (Friedkin, 1991; Saxena and Iyengar, 2020).

Degree centrality measures the connectivity of the network (how many edges/links per node). Vertices having neighbors with a higher degree centrality have more influence or information flow than those with lower degree centrality. Eigenvector centrality indicates the sum of the weights/scores of its neighbors' degree centrality. The closeness centrality is a measure of the average shortest distance from a vertex to each other vertex.

The betweenness centrality (BC) of a node is the number of shortest paths between any pair of vertices passing through that node. Nodes with high BC are gatekeepers controlling information flow dynamics and influence passing between others. It is hence a measure of central node(s) in between distinct clusters or communities within a nested complex network (i.e., modularity). Although a vertex can have a low degree centrality, and appear to be sparsely connected to the network, it can have a great influence on the whole network due to a high BC. Thus, BC is an emergent network feature in complex networks indicating the presence of modularity and recursiveness or nested community structures (i.e., hierarchical clusters). The Newman-Girvan modularity algorithm uses betweenness as a measure of community structure detection.

| Signaling Protein | Category | Brief Description of Key Functions |
|---|---|---|
| Complement | Immune Response | Major defense system of innate immunity. |
| ANNEXIN | Immune Response | Glucocorticoid-regulated group of proteins involved in multiple cellular processes. |
| MIF | Immune Response | Cytokine and key regulator of innate immunity and related processes such as macrophage migration. |
| GRN | Immune Response | Granulin plays role in developmental processes, inflammation, and cell proliferation. |
| LIGHT | Immune Response | Member of the TNF ligand family involved in multiple cellular processes including immune homeostasis. |
| TNF | Immune Response | Inflammatory cytokine regulating immune signaling. Activates the transcription factor NF-kB. |
| IL6 | Immune Response | Pro and anti-inflammatory Cytokine linked to TNFα expression. |
| IL10 | Immune Response | Cytokine regulating immune-inflammatory pathways and JAK-STAT signaling pathway. |
| CX3C | Immune Response | Chemokine mediating cell migration and adhesion of leukocytes to inflammed tissues/tumors. |
| VISFATIN | Immune Response | Adipokine (a type of cytokine) involved in glucose metabolism and inflammatory responses. |
| VEGF | Growth Factor | Angiogenesis and tumor microenvironment remodelling/cancer stem cell niche maintenance. |
| PDGF | Growth Factor | Key signal in glial cells and glioma cell growth, division, and differentiation dynamics. |
| ncWNT | Growth Factor | Complex processes including cell patterning/morphogenesis, development, and cell communication. |
| FGF | Growth Factor | Mitogenic/cell growth and differentiation. |
| TGFb | Growth Factor | Multifunctional cytokine involved in complex cancer processes including phenotypic transitions, metastasis, and cancer stem cell dynamics. |
| BMP | Growth Factor | Signaling molecules belonging to TGF-β superfamily involved in morphogenesis and differentiation. |
| GDF | Growth Factor | Growth differentiation factors belong to the TGF-β superfamily involved in developmental processes. |
| IGF | Growth Factor | Simulated by endocrine signals. Key regulator of |
| MK | Growth Factor | Also known as NEGF2, Heparin-binding growth factor involved in diverse processes such as tumorigenesis, and cancer stem cell renewal. |
| PTN | Growth Factor | Belongs to the same family of growth factors as MK and a cancer biomarker. |
| ANGPTL | Growth Factor | Secreted proteins structurally similar to ANGPT involved in inflammation, hematopoiesis, metabolism, and cancer processes. |
| ANGPT | Growth Factor | Vascular growth factor like VEGF. |
| SEMA3 | Neurotrophic | Related to neural system development and axonal guidance. Regulated by Ras/Rho signaling. |
| PSAP | Neurotrophic | Glycoprotein involved in Glioblastoma invasion and epithelial-mesenchymal transition/phenotypic plasticity. |
| PROS | Communication | Vitamin-K dependent blood clotting factor. |
| GALECTIN | Communication | Class of signaling proteins related to cell-cell interactions, cell-matrix adhesion and glycosylation. |
| COL | Communication | Collagen - extracellular matrix. Key component of cancer microenvironment mediating cancer adhesion, metastasis, and communication. |
| GAS | Communication | Tumor suppressor involved in multiple cellular processes including cell cycle regulation. |

**Table S1. Summary of key signaling proteins and functional pathways identified by the CellChat algorithm.** The role and classification of signaling proteins identified in Figure 4C-D in the Main Text.
